# Coordination of flux-related leaf and root traits impacts forest community assembly

**DOI:** 10.1101/2024.11.25.625305

**Authors:** Zeqing Ma, Mengcheng Duan, Lawren Sack, Chengen Ma, Le Li, M. Luke McCormack, Mingzhen Lu, Dali Guo

**Affiliations:** Qianyanzhou Ecological Research Station, Key Laboratory of Ecosystem Network Observation and Modeling, Institute of Geographic Sciences and Natural Resources Research, Chinese Academy of Sciences, Beijing 100101, China; Department of Ecology and Evolutionary Biology, University of California Los Angeles; Los Angeles, California, 90095, USA; The Center for Tree Science, The Morton Arboretum, Lisle, Illinois, 60532, USA; Department of Environmental Studies, New York University, New York City, New York 10003, USA; University of Chinese Academy of Sciences, Beijing 100049, China

**Keywords:** Above- and belowground, plant functional traits, leaf vein density, ecological strategy, hydraulics, root, woody angiosperms, community composition

## Abstract

Understanding the functions and coordination of plant traits is critical for predicting how diverse species respond to climate change. According to hydraulic and economic theories, leaves and roots—key organs for resource acquisition—are expected to function in coordination, such that species with faster resource utilization would possess leaf and root traits that facilitate rapid carbon, nutrient, and water uptake and fluxes. However, there has been limited evidence supporting leaf-root trait coordination and a role for that coordination on community structure. Here, we measured 13 leaf and root functional traits for 101 woody species from six tropical and subtropical forests, and assessed coordination and its association with community dominance. Hydraulic traits, such as leaf vein density and root vessel density, were coordinated between organs and showed compensation trade-offs between traits within organs, such as, leaf vein density and diameter. Economic traits relating to composition, such as nitrogen concentration, were coordinated between organs, whereas economic structural traits were decoupled, such as leaf mass per area and specific root length. Overall, hydraulic traits and economics traits were partially independent. The coordination of flux-related leaf and root traits was associated with ectomycorrhizal symbiosis and with dominance within the community. These findings indicate how trait organization within and across organs contributes to optimal whole plant function, with implications for performance in natural communities.

## Main text

Leaves and roots are the organs through which plants acquire light energy, carbon, water, and nutrients, the fuel and material for the growth of individuals and ecosystems. As the two key resource-acquiring organs, their coordination for resource uptake has been predicted in hydraulic and economic theories (Tyree & Zimmermann 2002; Reich 2014; Weemstra *et al*. 2023). However, evidence of the functional coordination of leaves and roots based on associations between their traits has been scarce and controversial (Bergmann *et al*. 2017; Carmona *et al*. 2021; Weigelt *et al*. 2021). Thus, extensive research has demonstrated tissue nitrogen (N) correlations between roots and leaves (Tjoelker *et al*. 2005; Westoby & Wright 2006; Geng *et al*. 2014; Weigelt *et al*. 2023). However, controversies persist regarding how leaf and root structural and hydraulic traits may be coordinated (Withington *et al*. 2006; Valverde-Barrantes *et al*. 2017; McCulloh *et al*. 2019; Weigelt *et al*. 2021; Verslues *et al*. 2022).

Theories for the coordinated function of leaves and roots assume that both organs are acted on by the same selective forces during evolution (Wright *et al*. 2004; Boyce 2005; Valverde-Barrantes *et al*. 2017; Weigelt *et al*. 2023). In that case, leaf and root traits might be unified along a single overall dimension of plant design (Freschet *et al*. 2010; Weigelt *et al*. 2023). The plant economic spectrum (PES) predicts that species with rapid resource uptake should achieve this with acquisitive economic traits in both leaves and roots, resulting in correlations between key traits including leaf mass per area (LMA) and specific root length (SRL) (Freschet *et al*. 2010; Reich 2014; Bergmann *et al*. 2020; Weigelt *et al*. 2021). Plant hydraulic theory also predicts that maximum flux rates would be coordinated across organs, as increasing transport through one organ alone would increase bottlenecks in other organs (Tyree & Zimmermann 2002; Noblin *et al*. 2008; Wolfe *et al*. 2023). These theories of unified selection to optimize leaf and root function would predict positive coordination of leaf and root hydraulic traits such as leaf vein density (i.e., vein length per unit area, VLA) and root vessel density (VesDensroot) (Fig. 1a), and potential “compensatory” trade-offs among functional traits within organs due to selection and/or allocation or design conflicts(Agrawal *et al*. 2010; Fox 2011; Garland *et al*. 2022), for example, VLA *vs.* leaf vein diameter (VeinDiamleaf) (Feild & Brodribb 2013).

**Figure 1.**
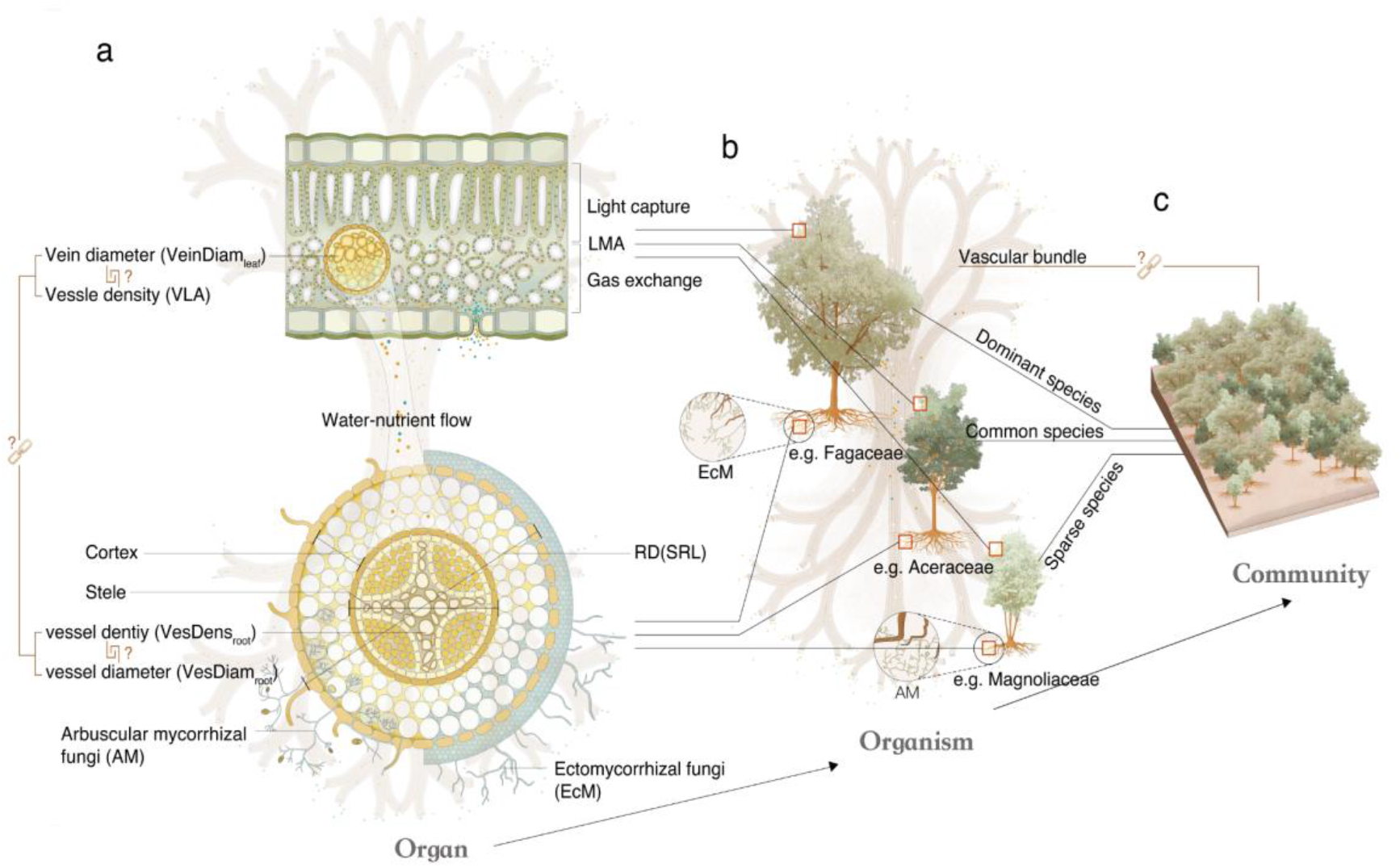
The coordination of root and leaf traits across scales from tissues within organs, to individual species performance to community assembly. **a,** At the organ scale, plant leaves are designed for efficient light capture and gas/water exchange, while plant roots and their associated mycorrhizal fungi support nutrient and water uptake. Water and nutrients enter through roots and are transported to plant leaves via the vascular system that coordinates whole-plant function. We hypothesized that the vascular hydraulic flux-related traits are coordinated between roots and leaves across species. Within organs, certain traits may show compensation trade-offs, such as vein density (VLA) and vein diameter (Veindiamleaf)(Feild & Brodribb 2013). b, Yet, each species will show a unique trait coordination, which would influence its performance in its community. For example, species of the family Magnoliaceae generally have thick roots with wide root vessels, and thick leaf minor veins, relative to other species, corresponding to a low abundance value in their home community. c, We hypothesized that across species, the opitmzied coordination between leaves and roots of vascular flux-related traits, and of economics-related traits, would be linked with mycorrhizal association and with their relative dominance within forest communities.

An alternative hypothesis is that roots and leaf traits are decoupled in response to distinct constraints and selective pressures (Weemstra *et al*. 2016; Carmona *et al*. 2021; Vleminckx *et al*. 2021). Even under the hypothesis of coordinated leaf and root function at whole-plant scale, their trait decoupling could arise if, independently of their structures (i.e., their quality), the amounts of leaves and roots (i.e., their quantity) vary widely. For example, a root system constructed of roots with relatively slow uptake rates can still achieve very high total system uptake if there are many individual roots. If so, the goal of understanding whole-plant organization may be considerably more complex, and depend, in part, on constraints on allocation to above and below ground parts (Freschet *et al*. 2015; Vleminckx *et al*. 2021; Weemstra *et al*. 2023).

Root-leaf relations are further complicated by trade-offs and interactions between roots and symbiotic fungi (Kong *et al*. 2019; Bergmann *et al*. 2020; Weigelt *et al*. 2021). Almost all trees form symbioses with arbuscular mycorrhizal (AM) or ectomycorrhizal (EcM) fungi to acquire nutrients and water against the background of competition with numerous soil microbes (Brundrett 2002; Liu *et al*. 2015; Chen *et al*. 2016; Meng *et al*. 2023). The great variety in the identities and intensities of mycorrhizal associations raises the fundamental question of how mycorrhizal fungi affect whole plant water and nutrient acquisition and transport efficiency, and thus, species competitiveness and community assembly (Fig. 1b). Notably, community ecology has historically emphasized niche differentiation according to leaf or stem traits, with much less focus on below-ground traits (Laughlin 2014; Laughlin *et al*. 2021; Da *et al*. 2023; Werden *et al*. 2023). A holistic understanding of whole-plant coordination and community organization will improve the prediction of species dynamics with different below and above -ground strategies, especially as species and ecosystems turn over under climate change.

Here, we focused on the coordination of root and leaf traits, association with the two main mycorrhizal types, and their scaled influence on the dominance of individual species within their respective communities. We first tested the generality of the plant economic spectrum, in which, across species, the leaf and root economic traits (LMA, leaf nitrogen (LeafN) vs. SRL, root nitrogen (RootN)) would show coordinated variation for rapid metabolism and growth, on the one hand, or slower, more conservative growth. Further, we hypothesized that across species, the vascular hydraulic transport-related traits would be coordinated between roots and leaves, with some traits showing compensatory trade-offs within organs. We especially focused on traits related to vascular hydraulic transport in roots and leaves, that is, the ratio of root stele to diameter (Stele:Diam), root vessel diameter and density, and leaf vein diameter and density. Despite species differing dramatically in their root and leaf design within a similar forest – each species might have its unique coordination (Fig. 1b) resulting in performance differences in its environment and community. We thus hypothesized that the optimized flux-related traits and mycorrhizal associations of individual species would scale up to an association with their relative dominance in forest community structure (Fig. 1c). Such a mechanism would assume a trait-by- competitiveness pattern(Bennett *et al*. 2016), in which a plant species optimizes coordination of different resource acquisition traits that can enhance its dominance in the community.

We measured 13 leaf and root traits for species with two contrasting mycorrhizal associations across 101 diverse woody angiosperms (59 genera within 31 families) from six subtropical and tropical forests (Table 1). This dataset is unique in that we: (i) applied consistent methods to measure traits, differing from database meta-analyses that used various sources and measurement methods; (ii) included hydraulic traits, such as root anatomy, often ignored in meta-analyses; (iii) focused on the first-order roots (the most distal absorptive fine roots), which are most comparable in function to leaves; and (iv) calculated an “importance value” for each species as a quantitative indicator for evaluating the species dominance in their natural community.

**Table 1.**
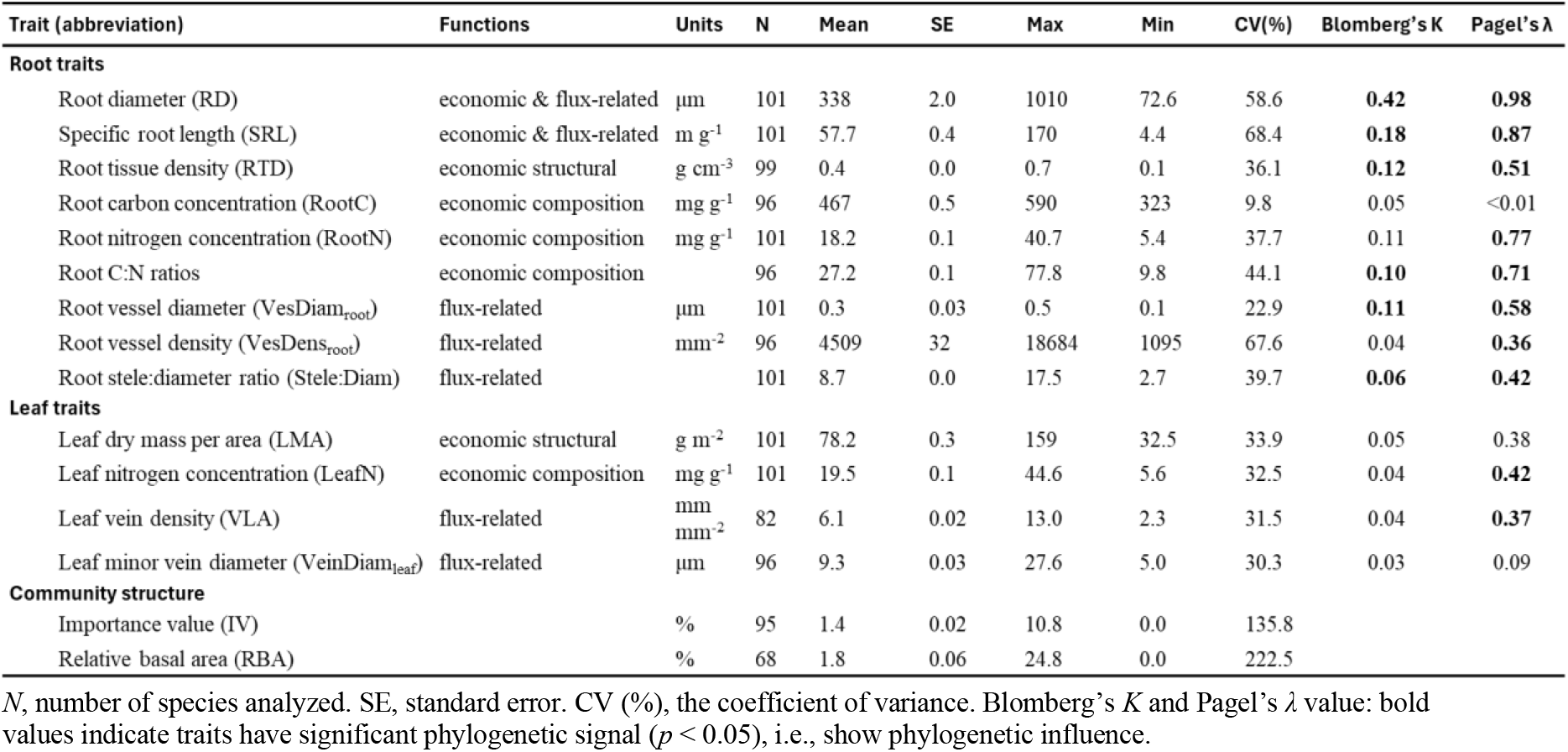
Summary of the 13 key leaf and root traits and species importance value for woody species in six (sub)-tropical forest.

## Results and Discussion

### Separation of flux-related and economic trait axes in roots and leaves

Within the (sub)-tropical woody plant trait space, we identified two main independent dimensions of variation (Fig. 2a; Extended Data Table. 4,5,6). Six root and leaf hydraulic traits (including SRL) were strongly loaded in the first axis, which explained 40.5% of the total variation, whereas economic traits, including leaf nitrogen (LeafN) and root nitrogen (RootN) and leaf dry mass per unit area (LMA), were loaded in the second axis (21.6%), indicating the decoupling of flux traits and economics traits.

### Between-organ coordination (and within-organ compensation) of flux-related traits in leaves and roots

Across species, the vein length per leaf area (VLA, i.e., leaf vein density) was positively related to root stele: diam ratio (Fig. 2b; *r* = 0.47, *P* < 0.001) and specific root length (SRL) (Fig. 2c; *r* = 0.30, *P* < 0.01), indicating coordinated vascularization in roots and leaves. These results support the expectation that rapid leaf transpiration is coupled with high capacity for root water delivery to maintain plant transport by optimizing traits that would minimize resistance to water flow in these organs that represent bottlenecks within the plant (Tyree & Sperry 1989; Steudle & Peterson 1998; Sack & Holbrook 2006; Verslues *et al*. 2022). Thus, root-leaf hydraulic flux coordination likely resulted from maintaining a continuous water potential gradient due to the continuity of the xylem.

**Figure 2.**
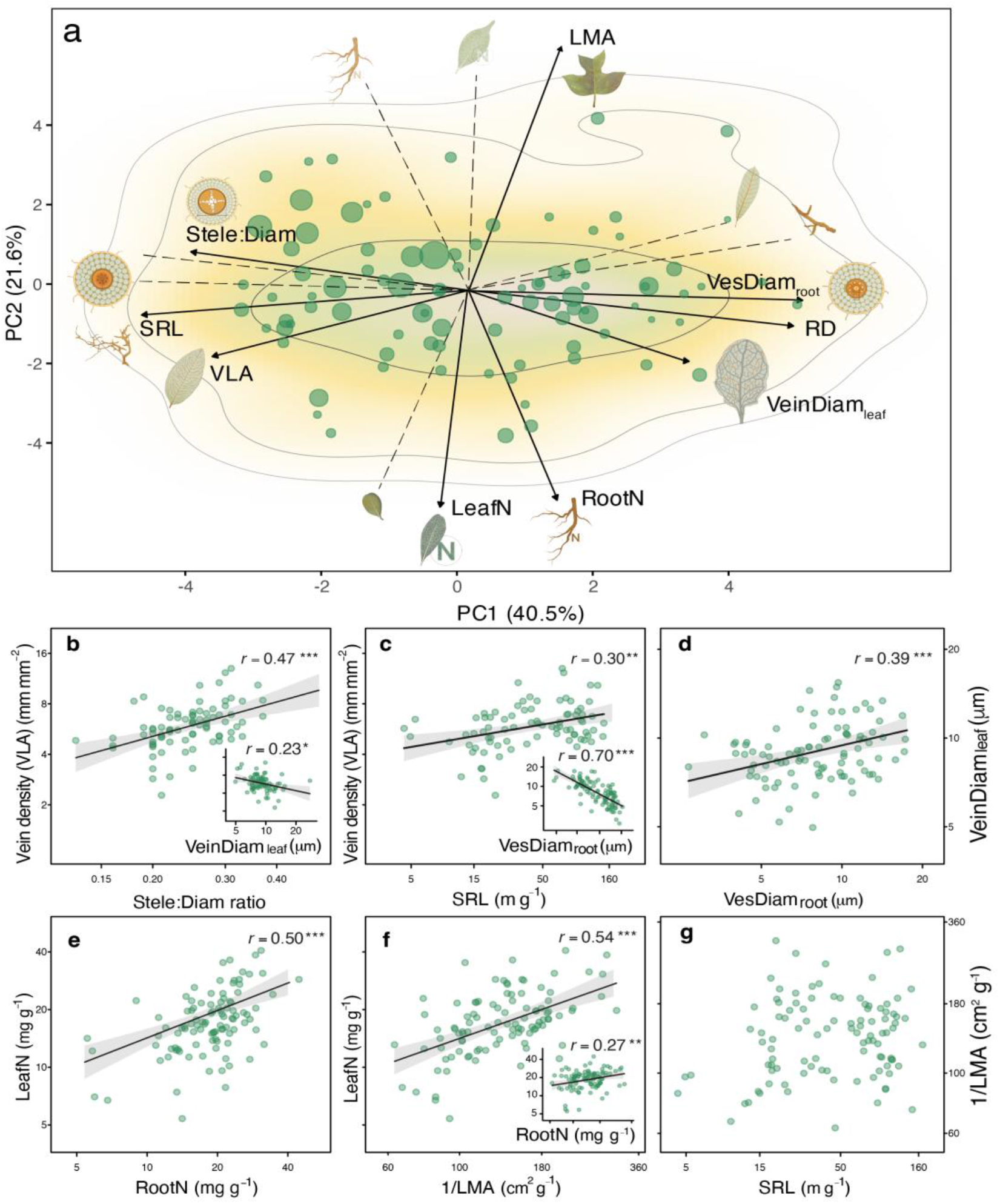
Root-leaf trait coordination and compensation across 101 subtropical/tropical forest species. **a,** Principal component analysis highlights independent axes of vascular-related traits (PC1) and economic traits (PC2) across 101 woody species in six (sub)-tropical forests. These two orthogonal axes suggest that tradeoffs operate independently along the hydraulic and economic dimensions. PC1 indicates significant correlations among root diameter (RD), Stele:Diam ratio, root vessel diameter (VesDiamroot), specific root length (SRL), vein length per unit leaf area (VLA), and minor leaf vein diameter (VeinDiamleaf). PC2 consists of the “economics traits”, showing strong correlations between root nitrogen (RootN) and leaf nitrogen concentration (LeafN), and leaf dry mass per area (LMA). We acquired importance values (relative coverage, relative abundance, relative dominance) for 95 species in the different native communities. Each species’ importance value is mapped onto a size scale; bigger circles associated with higher importance value. The color gradient in **a** indicates regions of highest (green) to lowest (white) occurrence probability of species in the trait space, with contour lines indicating 0.5, 0.95, and 0.99 quantiles. **b**, Stele:Diam ratio was positively related to VLA. **c**, SRL (proxy of root conduits density) of first-order roots is positively related to VLA (proxy for leaf conduits density) at the organ level. Both measures of conduit density, SRL and VLA, are key determinants of transport capacity. As the rules of the branching systems, thicker veins and root vessels come at the cost of their length density (inset panels, compensation in VLA *vs.* VeinDiamleaf, SRL *vs*. VesDiamroot), which is important for determining their transport efficiency. **d,** At the terminal ends of the distribution system, there is a positive coordination between VeinDiamleaf and VesDiamroot. The density of conduits (**b, c**) more than the size of conduits (**d**) is the basis of differences in transport in root or leaf, despite the long-standing expectation that the size of conduits is the leading determinant of transport capacity. Economic composition traits show between organ coordination (RootN *vs.* LeafN; LeafN *vs.* 1/LMA; RootN *vs.* 1/LMA) (**e**, **f**), but economic structural traits show decoupling between organs (**g**).

Across species, leaf vein diameter (VeinDiamleaf ) was positively related to the root vessel diameter (VesDiamroot) (Fig. 2d; *r* =0.39, *P*<0.001), indicating the xylem cell size coordination within whole-plant design. This cell size coordination also supports the conservation of hydraulic architecture during evolution (McCulloh *et al*. 2019; Koçillari *et al*. 2021) compared to root and leaf morphology. Indeed, we found that size-related root and leaf hydraulic traits showed a narrow range of variation, suggesting significant phylogenetic conservatism in conduits diameter (Table 1). Still, small increases in conduit diameter would disproportionately increase hydraulic conductivity, as the flow rate in capillaries is proportional to the fourth power of the radius as described by the Hagen–Poiseuille law (Tyree & Ewers 1991). The water entering plants is determined by two root parts—the stele and cortex (Steudle & Peterson 1998; Robinson *et al*. 2003). Generally, the water potential in the cortex would be close to that of soil water while stele water potential would be closer to leaf water potential (Robinson *et al*. 2003). The coordination of leaf and root hydraulic traits would thus contribute to a conservative gradient in water potential from the root stele to the leaf vein due to the anatomical continuity of the xylem. Within the hydraulic dimension, the absorptive roots serve as the water inlet and leaf stomata act as a water outlet.

We found evidence for compensation trade-offs within organs that are associated with whole-plant hydraulic coordination across organs (Fig. 2b-c, inset). Thus, root vessel diameter (VesDiamroot) was negatively related to SRL, and leaf minor vein diameter (VeinDiamleaf) was negatively related to vein density (VLA) (Fig. 2b-c, inset; Extended Data Fig 1). Smaller root vessel diameter and smaller leaf veins may be an adaptation to enable the development of a greater density of conduits (i.e., VLA, SRL), potentially at a reduced cost in space and materials (Sack *et al*. 2012). For example, the leaf vein diameter is typically negatively related to vein density (Feild & Brodribb 2013). This organization may help maximize the conduit network transport distribution efficiency with respect to construction costs. Notably, the size of conduits and the diameter of fine roots and veins are not the only determinants of transport capacity. Indeed, i) conduit number rather than diameter is the basis of differences in transport in these fine structures (Koçillari *et al*. 2021); ii) thicker veins and roots come at the cost of their length density (i.e., VLA, SRL) (Fig. 2; Extended Data Fig. 2), which is more important for determining their transport efficiency, and iii) at the terminal ends of the collecting or distribution system, most transport is radial, not axial. Therefore, for both leaf minor veins and absorptive fine root the exchange surface area would tend to be more important than the pipe diameters inside.

The coordinated evolution of root and leaf traits was supported by our phylogenetic analyses (Boyce *et al*. 2009). Most species in Magnoliaceae and Lauraceae tended to have thicker roots and wider root vessels, distinct from species of other more recently diverged families (Table.1, Extended Data Fig. 3, 7) (Kong *et al*. 2014; Ma *et al*. 2018). As atmospheric CO2 has decreased since the late Cretaceous, stomata density and vein density significantly increased to improve leaf carbon fixation (Raven 2002; Boyce *et al*. 2009; Comas *et al*. 2012). Given the increased water consumption, plants would also need to adapt to increase water acquisition (Boyce 2005; Boyce *et al*. 2009; Scoffoni *et al*. 2017). Thus, thinner root diameter and cortex would enable greater specific root length and root length density to optimize water acquisition (Rieger 1999; Hernandez *et al*. 2010; Masumoto *et al*. 2022). The selective force for increased CO2 uptake would drive the evolution of the hydraulic network from leaves to roots for maximized water flow (Fig. 2), leading to hydraulic coordination to a higher degree as the whole plant functions. As a result of these evolutionary processes, the small but dense leaf vein and root conduits enhanced the adaptive capacity to utilize pulsed resource supplies and helped plants forage for water and nutrients (Comas *et al*. 2012).

### Coordination and decoupling of economic traits in roots and leaves

Leaves and roots showed coordination in economic composition traits, i.e., in leaf (LeafN) and root nitrogen concentrations (RootN); LeafN *vs.* 1/LMA; RootN *vs.* 1/LMA (Fig. 2e, f). A leaf with higher nitrogen concentration and photosynthetic rates would match a root with a higher nitrogen and potentially higher respiration rate and nitrogen uptake rate (Volder *et al*. 2005), according to stoichiometric conservatism in active metabolic tissues (Sterner & Elser 2002). In addition, some nitrogen-rich proteins act as signal material or enzymes, enhancing coordination (Reich *et al*. 2008). Moderate positive coordination in nitrogen between leaf and root seems to be a general pattern globally (Reich *et al*. 2008; Wang *et al*. 2022), with variation primarily associated with soil substrate and temperature (Reich & Oleksyn 2004). The positive correlation in nitrogen suggests similar metabolism in the leaf and absorptive roots (Extended Data Fig.1, Fig. 2e), and integration of whole-plant functioning.

Notably, root economic structural traits — specific root length (SRL) and root diameter (RD) were strongly loaded at the first axis, while leaf mass per area (LMA) was strongly loaded at the second axis (Fig 2a; Extended Data Table 6), indicating that the economic structural dimension is not strongly coordinated between leaves and roots (Kramer-Walter *et al*. 2016; Weemstra *et al*. 2016; Han *et al*. 2023). Overall, ‘fast’ leaves with low LMA were not necessarily associated with high SRL for quickly acquiring soil resources (Liu *et al*. 2010; Holdaway *et al*. 2011; Fort *et al*. 2013; Poorter & Ryser 2015; Weigelt *et al*. 2021). One of the main reasons for this decoupling is that leaves and roots can adjust biomass allocation to maintain the functional balance (Poorter *et al*. 2012; Freschet *et al*. 2015; Freschet *et al*. 2021).

Further, structurally, the two organs have fundamentally distinct designs (Vogelmann *et al*. 1996; Brodribb *et al*. 2010), with leaves generally deploying a two-layer laminar design (Fig. 1a), where the upper mesophyll cells (palisade and spongy) layer primarily capturing the light and radiant energy, and the lower epidermis and stomata maximizing the CO2 uptake and minimizing water loss (Brodribb *et al*. 2007; Sack & Scoffoni 2013). These two layers may vary independently (Li *et al*., 2015), leading to the same LMA but different functional modules (Li *et al*., 2017). By contrast, roots follow a multi-layer cylinder design (Robinson *et al*. 2003; Guo *et al*. 2008), and the cortex and stele tightly co-vary (Kong *et al*. 2014). Despite the general decoupling of the leaf and root structure owing to their unique design requirement, we infer that the physical structure of the leaf layer containing flux-related parts might coordinate with root anatomical traits owing to their hydraulic connectivity.

Distinct resource types may also facilitate decoupling of economics modules. Roots are designed differently from leaves to acquire physical mass resources rather than light energy and air flux. Water and nutrients require physical connections to soil, and all mass must cross the epidermis, cortex, and endodermis before entering the xylem (Fig. 1a) (Steudle & Peterson 1998; Barlow 2002). Leaves have diverse shapes, functioning as semi-closed systems protected by various tissues (Fig. 1a), and demonstrate multi-functionality, finely tuning their behavior to manage water loss, light capture, and carbon fixation (Westoby *et al*. 2002; Laughlin 2014; Díaz *et al*. 2016). The diverse shapes and complex functions of leaves make them difficult to compare to the simple cylindrical shape of roots. Overall, root and leaf morphological traits are independently adapted and compete either belowground for heterogeneous soil resources or aboveground for homogeneous CO2, respectively (Boyce 2005; Freschet *et al*. 2015).

Overall, we found that flux-related traits were decoupled from economics traits. Specifically, hydraulics traits display between-organ coordination and within-organ compensation tradeoffs while economic composition traits indicate between-organ coordination but economic structural traits suggest decoupling between organs (Fig. 2).

### Mycorrhizal association mediates root-leaf functional coordination

We investigated the degree that the coordination patterns between leaf and root traits might differ with mycorrhizal type. Thus, we tested these patterns separately for the AM- and EcM- plant species (Extended Data Table 7), which are thought to support distinct strategies for resource acquisition belowground and nutrient cycling (Phillips *et al*. 2013).

The correlation among certain hydraulic traits (Stele:Diam *vs.* VLA) and among economic composition flux traits (RootN *vs.* LeafN) were significant across both mycorrhizal types. However, for given trait associations, the patterns were found between AM- and EcM species (Extended Data Fig. 2; Table 7). It is likely that, in these cases of pairwise relationships, the correlation between traits were mediated by mycorrhizal type (Extended Data Table 8). Thus, species with thick roots can employ mycorrhizal hyphae to achieve high absorptive surface area equivalent to that of species with much thinner roots (Eissenstat *et al*. 2015; Liu *et al*. 2015), essentially achieving functional equivalence (e.g., total absorptive surface) albeit with quite distinct root morphological traits (McCormack & Iversen 2019). This influence of mycorrhizal association can potentially complicate the root-leaf trait coordination, which might help explain the variations we observed (Extended Data Fig. 2).

### Flux-related root-leaf traits influence species dominance within a community

The discovery of root-leaf coordination in water flux-related traits raises the question of whether such coordination might impact species dominance within a community setting (Fig. 1; Fig. 3; Extended Data Fig. 4). Further, given the close relationship between roots and their mycorrhizal partners, it will be important to also consider how these relationships contribute to tree species dominance. The leading hydraulic flux-related dimension (PC1, mainly driven by SRL, Stele:Diam ratio, VesDiamroot and VLA) was associated with tree species importance value (Fig. 3a; *r* = 0.32, *P* < 0.01) and relative basal area (Extended Data Fig. 4a) within diverse forest communities.

**Figure 3.**
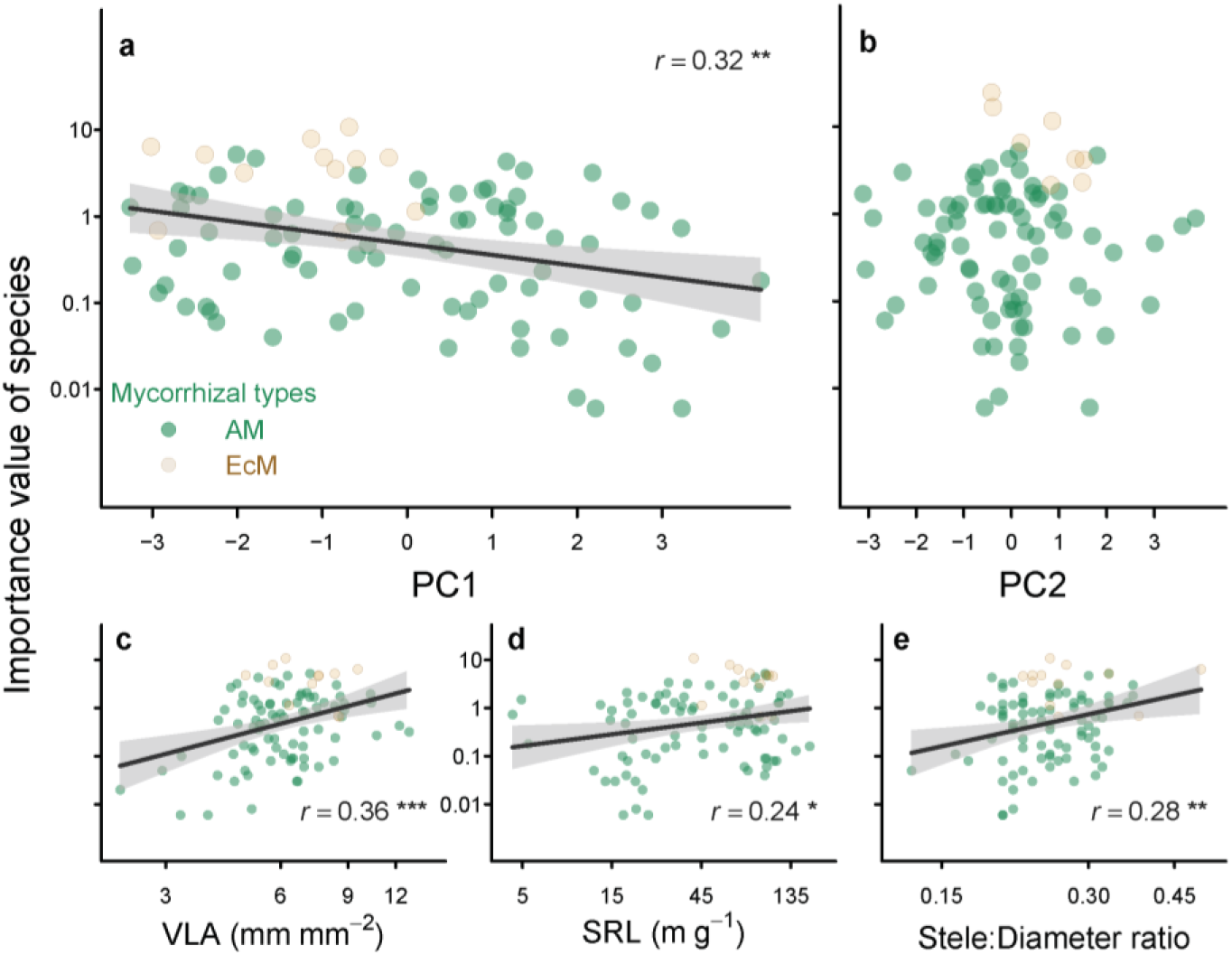
Flux-related root-leaf traits link to the species dominance in the communities. Different species with multiple water-use and light-capture strategies among different mycorrhizal types each occupy a unique position within their respective communities, ranging from low to high dominance indicated by their importance value (IV). **a,** we found a correlation between a species position within PC1 and their importance value (*R*^2^ = 0.09, *P* < 0.01). PC1 is the linear combination of a range of important hydraulic traits, with low values of PC1 denoting species with low root vessel diameter, low leaf vein diameter, low root diameter and high SRL (proxy of root conduits density), high vein leaf per leaf area (proxy of leaf conduits density), high Stele: Diam (Fig. 2a). **b**, we found no correlation between PC2 and species importance value. PC2 is a linear combination of leaf mass per unit area (LMA), leaf nitrogen concentration (LeafN), and root nitrogen concentration (RootN). **c**, **d**, conduit number (i.e., VLA, SRL), indicated that the exchange surface area, rather than diameter, is the basis of differences in transport in these fine structures, significantly influencing the performance of species in the communities. **e**, Stele:Diam ratio is important for determining their transport efficiency, constrained by the phylogenetic history, and is correlated with the importance value of species in the communities. Yellow and green represent the mycorrhizal fungi type; the EcM species tend to competitively dominate in seasonal forests. The x and y-axis were log scaled.

This association suggests that species with enhanced hydraulic capacity may exploit larger resource hypervolumes, contributing to their ecological dominance. The individual traits SRL, Stele: Diam ratio, and VLA are also closely associated species importance value (Fig. 3c-e), whereas economic traits show little association (Extended Data Fig. 5).

Based on a linear mixed effects model, we found that the tree mycorrhizal identity had a great influence on the species importance value in a given community (Fig. 4), stronger than plant functional types such as growth form (tree vs shrub), leaf habit (evergreen vs deciduous) across 95 species (Extended Data Table 8, 9). We observed that EcM species had a higher importance value, on average, than AM species in these seasonal forests (*P* < 0.001, linear mixed-effects model). These EcM host species typically have higher leaf vein density and greater specific root length, also associated with their dominance in these (sub)-tropical forests. These findings suggest that symbioses-root-leaf coordination scales up as an influence on the evolutionary assembly of plant communities and plant–plant interactions.

**Figure 4.**
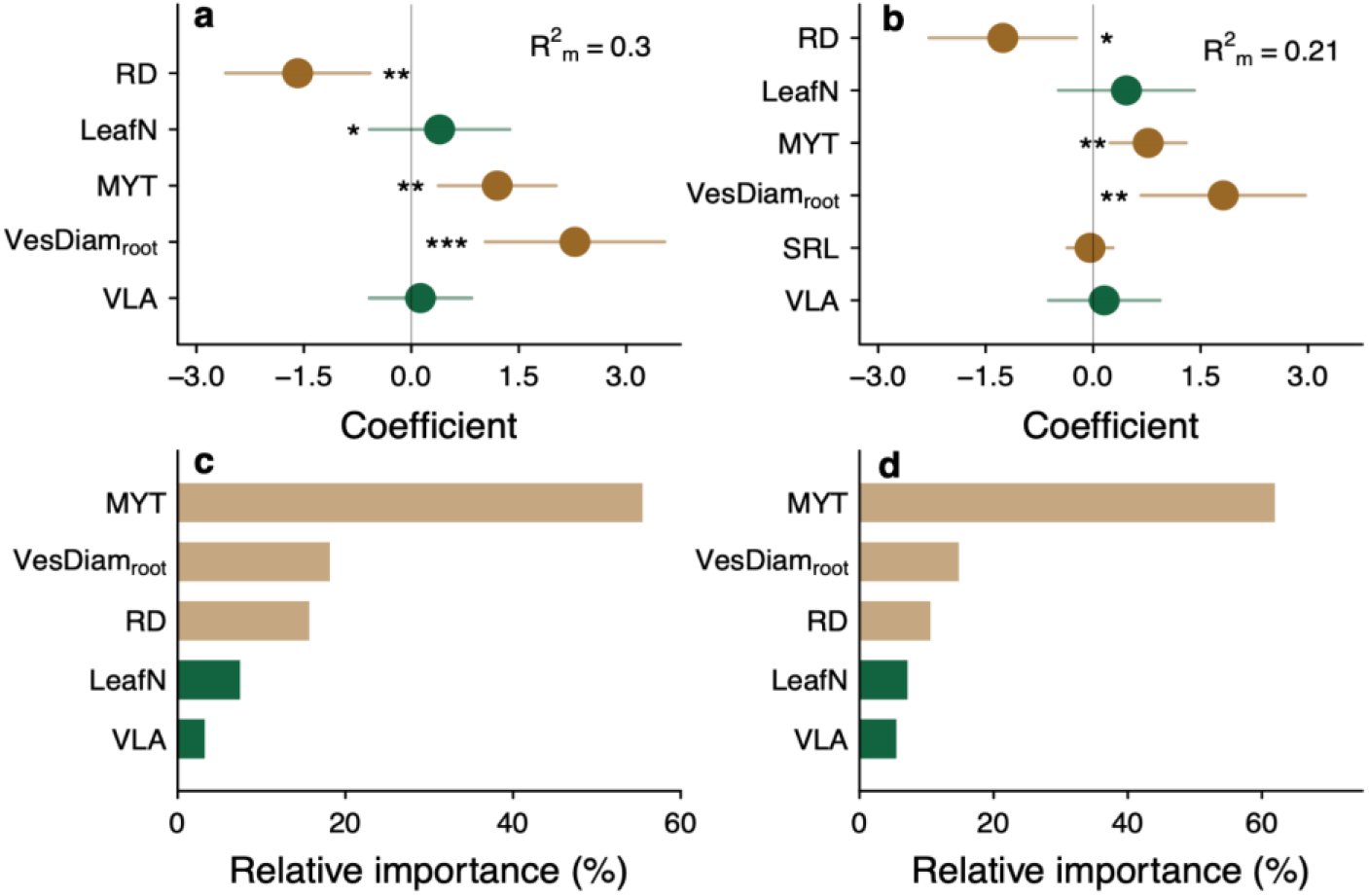
The influence of key root and leaf traits on the species’ importance value (IV). We performed linear mixed-effects models, with all 9 root-leaf traits and plant functional type (mycorrhizal associations, leaf habit and growth form) as fixed effects and species phylogeny and sampling sites as random effects. **a**, **b**, Optimal model results for predictors of species IV using the complete and imputed datasets, respectively. Root diameter (RD), root vessel diameter (VesDiamroot), and mycorrhizal type (MYT) were significantly correlated with the IV of species. We also interpreted the key factors of variation in the species IV in the community by variance partitioning, based on the complete (**c**) and imputed datasets (**d**). Root traits play a significant role in community assembly, and the relative importance of VesDiamroot, a flux-related trait, is four times greater than Leaf N concentration (LeafN). SRL: specific root length; VLA: leaf vein density.

There are at least three mechanisms for the advantages of EcM host species with thin roots and greater leaf vein density in the community. First, EcM trees with thin roots are typically more tolerant of water stress than AM trees in seasonal forests (Brzostek *et al*. 2014). Thin roots are relatively easily coated with EcM (Comas *et al*. 2014; Kong *et al*. 2019), increasing water foraging space with hydrophilic hyphae (Agerer 2006). Thin roots also may work as circuit breakers of the plant hydraulic system (McCulloh *et al*. 2019; Cuneo *et al*. 2021), and can quickly sense signals of soil drought (Bais *et al*. 2006), showing strong structural plasticity and flexibility to adapt to seasonal water supplies. As a result, EcM roots possibly can adjust better to pulsed water and wet-dry season shifts (Liese *et al*. 2019). Second, EcM may form a network of mycelium and some may decompose organic matter directly to obtain nutrients, enabling these associations to gain a nutritional advantage compared to their AM neighbors (Lindahl & Tunlid 2015). Third, the Hartig net of EcM protects the host roots from various pathogens and predators: this effect is especially important when species are in the early stage (i.e., seedlings) (Connell & Lowman 1989; Peay *et al*. 2010; Chen *et al*. 2019). As to their leaf design, EcM species in these forests typically have higher leaf vein density, higher hydraulic conductivity, and shorter vein spacing compared with AM species, which can make water transfer to mesophyll cells more localized, effectively supporting photosynthesis (Sack & Holbrook 2006; Brodribb *et al*. 2010; Feild & Brodribb 2013).

Overall, our results identify impacts of flux-related root-leaf traits and mycorrhizal association on plant performance and forest community assembly (Fig. 3, 4, Extended Data Fig. 6). Selective pressures in these seasonal forests may favor hydraulic designs that efficiently transport water, enable flexible nutrient foraging, and minimize construction costs. We speculate that the innovation of EcM association plus higher specific root length and leaf vein density might enhance the root resistance to herbivores (Chen *et al*. 2019)and plants’ adaptive ability to variable soil water supply, in turn, these plant species had higher survivorship during the long history of disturbances in these seasonal forests (Brzostek *et al*. 2014). Our trait-based evidence points to probable explanations for why EcM species and families (e.g., Dipterocarpaceae, Fagaceae, Myrtaceae) have the ability to become dominant species at (sub)-tropical latitude despite the exceptionally high diversity of AM tree species coexisting in these communities (McGuire *et al*. 2010; Peay *et al*. 2010; Lu & Hedin 2019).

By linking whole-plant traits organization with their influence on community dynamics, our study extends the current knowledge boundary of trait-based ecology. Instead of a “single economic dimension” design (Reich 2014), a plant may be constructed by different resource-capturing modules with multiple dimensions. Yet, all these dimensions are integrated by the whole-plant level vascular system to form a coherent above-belowground resource supply strategy. Further, our results point to the ability to use easier-to-measure leaf hydraulic traits to infer or predict root traits that are much harder to measure, as a foundation for fully parameterizing the hydraulic architecture of plants, a mission that is of high importance in the modeling ecosystems under climate change.

## Methods Summary

We collected leaf and root samples from 101 diverse woody angiosperms (59 genera within 31 families) from six (sub)-tropical forests, covering key clades of common species in southern China (Table 1; Extended Data Table 1, Fig. 6). We focused on mature plants and sampled at least three per species. Root samples were collected following the procedure described in Guo *et al*. 2008. We focused on the first-order roots, as they can be considered functionally comparable to leaves regarding resource acquisition. Leaf samples were collected from the upper part of the tree canopy, with > 40 mature fully expanded sun-exposed leaves collected from each individual trees.

We measured 13 key functional traits of leaves and absorptive roots associated with plant hydraulics, chemistry, and morphology (Table 1). The Methods provide more details of trait measurement*s*. We constructed a plant phylogeny using rbcL and matK sequences of each species (Extended Data Table 2). The phylogenetic tree was constructed using maximum likelihood and Bayesian approaches, with divergence time estimated by BEAST 1.7.1 (Drummond *et al*. 2012).

The importance value (IV) is an integrated measure of a species’ density, occurrence frequency, and basal area, offering a comprehensive assessment of its potential niche hyperspace and competitiveness within a community (Curtis & McIntosh 1951; Whittaker 1972). We obtained IV data for 95 of the 101 species examined, sourcing data from two primary sources. For 82 species, IV data was obtained from survey data collected in natural forest communities managed by the China Ecosystem Research Network (CERN), publicly available for approval through their website (http://www.cnern.org.cn). The remaining 13 species’ IV data were derived from published literature describing the community structure of nearby forests (Zeng *et al*. 1999; Fang 2005; Xu *et al*. 2015; Wen *et al*. 2018). The importance value for each species across different communities was calculated as the sum of relative density, relative frequency, and relative basal area (RBA), divided by three (Mueller-Dombois & Ellenberg 1974). The absence of IV for six species is likely due to their low abundance within these forest communities or the limited spatial scale of the community census. Further, we obtained the relative basal area (RBA), another indicator of species’ dominance, for 68 out of the 101 species from these data sources. Both IV and RBA were used to assess the influence of root-leaf traits on species’ roles in the community.

We performed trait imputation using Multivariate Imputation with Chained Equations (MICE) based on ecological and phylogenetic relationships between species to obtain complete trait information. The imputed datasets were compared with the complete dataset to evaluate the reliability of the imputation procedure (Extend Data Fig. 3, Table 4, 5, 6).

## Statistical analyses

All statistical analyses were performed using the R software (version 4.2.0.). Mean, minimum, maximum, and coefficient of variation (CV) were estimated for each trait across 101 species. Blomberg’s K and Pagel’s λ tests were conducted to assess the phylogenetic signal of each trait. Phylogenetically independent contrasts (PICs) were calculated to examine the effect of phylogeny on interspecific comparisons. Principal component analysis (PCA) was performed on transformed and standardized root-leaf traits of the complete (95 species) and imputed (101 species) datasets. Varimax rotation was applied to enhance interpretability. Phylogenetic PCA was also employed to reduce evolutionary noise and reveal functional coordination between root and leaf traits. Dissimilarities (Euclidean distance) among leaf habits, growth forms, mycorrhizal types were calculated using the permutational multivariate analysis of variance (PERMANOVA). The relationships between leaf and root traits were analyzed using standardized major axis regressions for all species and separately for AM and EcM species. A linear mixed model with restricted maximum likelihood was fitted to examine the impact of root-leaf traits and plant functional groups on species competitiveness. Species IV was used as the response variable, with nine key root-leaf traits, mycorrhizal types (AM vs. EcM), growth form (Tree vs. shrub), leaf habit (deciduous vs. evergreen) included as explanatory variables. Before fitting the model, we streamlined the fixed effects by selecting significant explanatory variables with a percent increase in mean squared error (%IncMSE) greater than zero using a random forest analysis (Liaw & Wiener 2001). We then used the dredge function in the MuMIn package (Barton 2009) for the model selection procedure.

## Materials and Methods

### Sampling approach

We collected leaf and root samples from the six tropical-subtropical forests of South China. The sites covered a latitudinal range from 18°40′ N to 24°32′N, with mean annual temperature ranging from 10.7 to 22.4 °C, and mean annual precipitation ranging from 1539 to 2651 mm (Extended Data Table 1). We sampled 101 angiosperm woody species (73 trees and 28 shrubs) from 70 genera and 33 families, covering key clades of common species in southern China, such as magnoliids, fabids, and lamiids (Extended Data Fig. 1b). All these species are native and dominant in the sub-canopy and canopy layers.

We sampled at least three mature individuals for each species to derive the mean species trait value. We collected root samples following the procedure described in Guo *et al*., (2008). We first removed the surface soil (0–20 cm) at the base of the sample trees to expose the main lateral roots. We selected and cut these root branches with intact terminal branch orders (> 5 g of fresh first-order roots to ensure accuracy of measurements). Subsamples of the roots from each tree were gently washed in deionized water to remove soil adhering to the roots. These samples were immediately put into plastic tubes filled with formalin-aceto-alcohol (FAA) solution (90 ml of 50% ethanol, 5 ml of 100% glacial acetic acid, and 5 ml of 37% methanol) for later anatomical measurements. The remaining samples were transported in plastic bags in a cooler to the laboratory and frozen at -20°C until subsequent morphological and chemical analyses.

We collected leaf samples from the upper part of the tree canopy by tree climbing, with > 40 mature fully expanded sun-exposed leaves collected from each individual trees. Once collected, more than five leaves per individual were immediately put into a buffered FAA fixation solution to analyze anatomical and venation traits. The remaining 15-25 leaf samples were used to measure morphological and chemical traits.

### Trait measurement

We measured fourteen key functional traits of leaves and absorptive roots (first-order root) that are thought to play an important role in resource acquisition and transportation in woody plants (Fig. 1; Table 1). We especially focused on the traits of first-order roots because these roots, which are the most short-lived and metabolically active, can be considered functionally comparable to leaves as resource acquisition organs (Pregitzer *et al*. 2002; Guo *et al*. 2008; Xia *et al*. 2010; McCormack *et al*. 2015). For anatomical traits, we randomly chose 20 first-order root segments fixed in FAA solution in the field from each species. The root segments were immersed in a sequence of alcohol solutions for dehydration before being embedded in paraffin (Guo *et al*. 2008). We sliced the roots into eight-μm-thick cross-sections. The cross-sections were then stained with safranine and fast green, with the cortex staining green and the stele staining red, and photographed using a Leica DFC450 camera (Nussloch, Germany) mounted on a Leica DM 2500 microscope. To ensure the maximum quality of the slices, we initially selected well over 20 root cross-sections and randomly chose 20 from all successful segments for our anatomical trait measurements. The anatomical traits (root diameter (RD), mean vessel diameter (VesDiamroot), stele diameter, and root vessel density (VesDensroot)) were measured using ImageJ (NIH Image, Bethesda, MD, USA). The ratio of stele diameter to root diameter (Stele:Diam) was calculated to indicate the proportion of resource transportation within a root.

More than five intact root branches for each species were dissected for morphological measurement as described in Pregitzer et al. (2002). The length of the first-order roots was measured using an electronic digital caliper. Specific root length (SRL) was calculated as the root length divided by its dry mass. The oven-dried (60°C, 48 h) root subsamples were ground to fine powder using a SPEX 8000-D mixer mill (SPEX, Edison, NJ, USA), and their N concentrations (RootN) were measured using an elemental analyzer (Vario Microcube; Elementar, Hanau, Germany).

For leaf venation measurements, paradermal sections were prepared according to the general protocols described by Carins Murphy *et al*. (2012); Brodribb *et al*. (2013), with modifications depending on the species. The adaxial epidermis and palisade mesophyll were carefully removed using a sharp razor blade to expose the minor veins. Sections were then placed in bleach (50 g L^-1^ sodium hypochlorite and 13 g L^-1^ sodium hydroxide) for several hours to several days, depending on the species, until clear. For some species that resisted clearing in bleach alone, sections were first placed in 5% KOH. After clearing, sections were carefully rinsed to remove bleach and stained in 1% toluidine blue. For each section, ten fields of view were photographed. Using ImageJ, leaf minor vein diameter (VeinDiamleaf, μm) was measured, and leaf vein density (VLA, mm mm^-2^) was calculated as the total vein length per unit leaf area. For each species, at least four leaves from different individuals were used, and 30 fields of view (each field with an area of 1250 ×937.5 μm) were selected between the midrib and the margin.

For leaf morphological and chemical traits, leaf surface area was scanned in the field site immediately after sampling using a portable scanner (Canon LiDE 110, Tokyo, Japan) and then measured using ImageJ software (NIH Image, Bethesda, MD, USA). The leaf samples were oven-dried to a constant mass at 60 °C for 48 hours. The dried mass of leaf samples was estimated with a precision of 0.1 mg. Leaf dry mass per unit area (LMA, g m^-2^) was calculated as the ratio of leaf dry mass to projected leaf area. The oven-dried leaf samples were ground to fine powder. Leaf carbon and nitrogen concentration (LeafN) was determined using an elemental analyzer (Vario EL III, Elementar, Hanau, Germany). We classified these traits as associated with plant hydraulics, chemistry, and morphology at the leaf and root (Table 1).

### The construction of plant phylogeny

Plant genomic DNA of each species was extracted from silica gel-dried leaves, which were collected at the same time as the roots were sampled. We determined each species’ rbcL and matK sequences (chloroplast gene fragments) (Extended Data Table 2). After model selection using jModelTest v2.1.1 (Posada 2008), the phylogenetic tree was constructed using maximum likelihood (ML) and Bayesian approaches. In the phylogenetic analyses, the tree branch length was proportional to the difference in divergence time between neighbor clades. Divergence time was estimated by BEAST1.7.1 (Drummond *et al*. 2012) with eight fossil calibration nodes ( Extended Data Table 3).

### Evaluation of species competitiveness

Importance value (IV) is a widely used measure of a species’ competitiveness within a natural community. We obtained IV data for 95 out of 101 species from nearby forest communities. To determine the IV for each species within its respective community, we collected community survey data from six long-term forest plots between 2015 and 2018. These plots are part of the Chinese Ecosystem Research Network (CERN) ecological stations (see Acknowledgements) and are subject to comprehensive forest inventories every five years. Our plant tissue sampling sites were located near these plots.

During the community survey, each forest plot was divided into multiple 10 ×10- meter survey units. In each survey plots, woody plants with a diameter at breast height (DBH) ≥ 1 cm were marked and identified, and their DBH was recorded. The data includes the basal area of each species, the number of stems for each species, the frequency of occurrence (the proportion of survey plots in which the species appears out of the total number of survey plots). The IV of each species was calculated as follows:

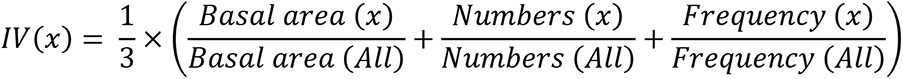

Where *x* is a particular species on a plot, and *All* is the sum of all species on each plot.

Relative basal area (RBA) is another indicator of species’ dominance. We obtained RBA data for 68 out of the 101 species from the same data sources. In assessing the impact of root-leaf traits on plant dominance, both IV and RBA serve as comparative benchmarks, helping to identify key root and leaf traits that influence community assembly.

### Plant functional group

Plant growth form (tree vs shrub), leaf habit (deciduous vs evergreen) and light demand (shade-tolerance vs sun preference) were provided in community census data. The identification of mycorrhizal types in all woody plants relies on the anatomical examination of absorptive roots. Plants characterized by a dense mycelial sheath encasing the root surface were categorized as ectomycorrhizal (EcM) species, while those with cortical cells penetrated by arbuscules were designated as arbuscular mycorrhizal (AM) species.

### Trait imputation

Our dataset encompassed nine root-leaf traits of 101 woody species in Chinese tropical-subtropical forests, with only 2.7% missing values. Notably, the data for all traits were nearly complete, with the sole exception being VLA, which had 19 missing values due to insufficient material available for measurement (Table 1). 82 species have complete trait measurements for the key nine traits forming the ‘complete dataset’. We employed Multivariate Imputation with Chained Equations (MICE) to obtain complete trait information, leveraging ecological and phylogenetic relationships between species. This method is chosen for its superior accuracy and reduced bias compared to single imputation methods (Cooke *et al*. 2019). The first ten phylogenetic eigenvectors were incorporated into the matrix to be imputed. To capture the uncertainty in the imputation process, 20 trait datasets were imputed and then averaged (‘imputed dataset’). These imputed datasets are based on the same input trait data but differ in their estimations for the missing data. Before the imputation process, all traits were log10-transformed. The reliability of the imputation procedure was evaluated by comparing statistical results for complete and imputed datasets.

## Statistical analysis

All statistical analyses were performed using the R software (4.2.0.). We estimated the mean value, minimum, maximum, and coefficient of variation (CV) for each trait across all 101 species (Table 1). We employed Blomberg’s K and Pagel’s λ test assuming a Brownian evolution model to test each trait’s phylogenetic signal. Phylogenetically independent contrasts (PICs) were subsequently calculated for all the functional traits to examine the effect of phylogeny on interspecific comparisons. Both descriptive statistics and phylogenetic analyses were performed using the original dataset, without incorporating imputed trait values derived from phylogenetic relationships.

A principal component analysis (PCA) was performed to examine the coordination of root-leaf traits using the transformed and standardized traits of the ‘complete dataset’ (82 species) and ‘imputed dataset’ (101 species), respectively. Two independent functional trait dimensions were identified via PCA. Subsequently, we applied a varimax rotation to the selected components to improve the results’ interpretability.

The phylogenetic PCA was also used to reduce the evolutionary noise, revealing the possible functional coordination between root and leaf traits under an evolutionary context.

For exploring the impact of different groups of species on functional trait variation, we grouped species according to their life habit (Deciduous vs. Evergreen), growth form (Tree vs. Shrub), mycorrhizal types (AM vs. EcM). The dissimilarities (Euclidean distance) among these groups of species were calculated along two independent trait variation axes using the permutational multivariate analysis of variance (PERMANOVA) (vegan package).

The relationships between leaf and root traits were analyzed by standardized major axis regressions (Warton *et al*. 2006). We first conducted this analysis across all species and then separately for AM and EcM species. In contrast to simple linear regression, we used SMA regressions because they do not assume a unidirectional effect of one parameter over the other (i.e., SMA minimizes the areas of the triangles formed by the observations and the line).

We fitted a linear mixed model with restricted maximum likelihood using species IV as the response variable to determine how root-leaf traits and associated plant functional groups affect community structure. As explanatory variables, we included nine key root-leaf traits, mycorrhizal types, growth form and leaf habit. To reflect our dataset’s large-scale spatial and phylogenetic structure, we treated the sampling sites and species phylogeny as crossed random effects. Before model fitting, we simplified the fixed effects by screening out the important explanatory variables with a percent increase in mean squared error (%IncMSE) larger than zero using a random forest.

We then standardized the response variable and all fixed effects to allow for a direct comparison of estimates. We fitted this simplified model with the R lme4qtl package using the lme4qtl function and diagnosed the normality and homogeneity of residuals (Ziyatdinov *et al*. 2018). To obtain the best-fitting models, we created a set of models with all possible combinations of the initial variables and sorted them according to the Akaike Information Criterion (AIC) fitted with Maximum Likelihood. We performed a model-averaging procedure based on the AICc (ΔAICc < 2) to determine parameter coefficients for the best final set of predictors of IV. This model selection procedure was performed using the dredge function in the MuMIn package. We ran linear regression between IV and root-leaf traits to test the effect of individual traits on IV.

## Data availability

All data will be deposited at Figshare upon publication of this paper.

## Supporting information

Supplementary files for root-leaf article

## Acknowledgments

We thank Shenglei Fu, Zhongliang Huang, Yuhong Liu, Yiping Zhang, Handong Wen, Xiaobao Deng and Mingxian Lin for their assistance in field sampling, Deliang Kong for assistance in data collection. We thank the following field research stations and government agencies for their kind support: Dinghushan and Heshan Station for Subtropical Forest Studies of South China Botany Garden, Chinese Academy of Sciences; Xianhu Botanic Park of Shenzhen; Ailaoshan Station for Subtropical Forest Studies and Xishuangbanna Station for Tropical Forest Studies of Xishuangbanna Tropical Botanical Garden, Chinese Academy of Sciences; Jianfengling Nature Reserve Management Bureau. This study was funded by the Natural Science Foundation of China (NSFC grant nos. 31325006 and 31530011).

## Author Contributions

ZM and DG developed the conceptual approach, compiled the data, and drafted the manuscript; LS, ZM, and ML developed the analytical approach, and all authors contributed to revisions. Conceptualization: ZM, DG, LS, ML; Methodology: ZM, DL, LS, ML; Investigation: CM, LL, DM; Visualization: ZM, ML; Funding acquisition: ZM, DG; Project administration: ZM, LS; Supervision: LS, ZM, DG; Writing – original draft: ZM; Writing – review & editing: ZM, LS, ML.

## Competing Interest Statement

The authors declare no competing interests.

**Classification:** Biological Sciences (Ecology)

