## Supplementary files for root-leaf article for "Coordination of flux-related leaf and root traits impacts forest community assembly"

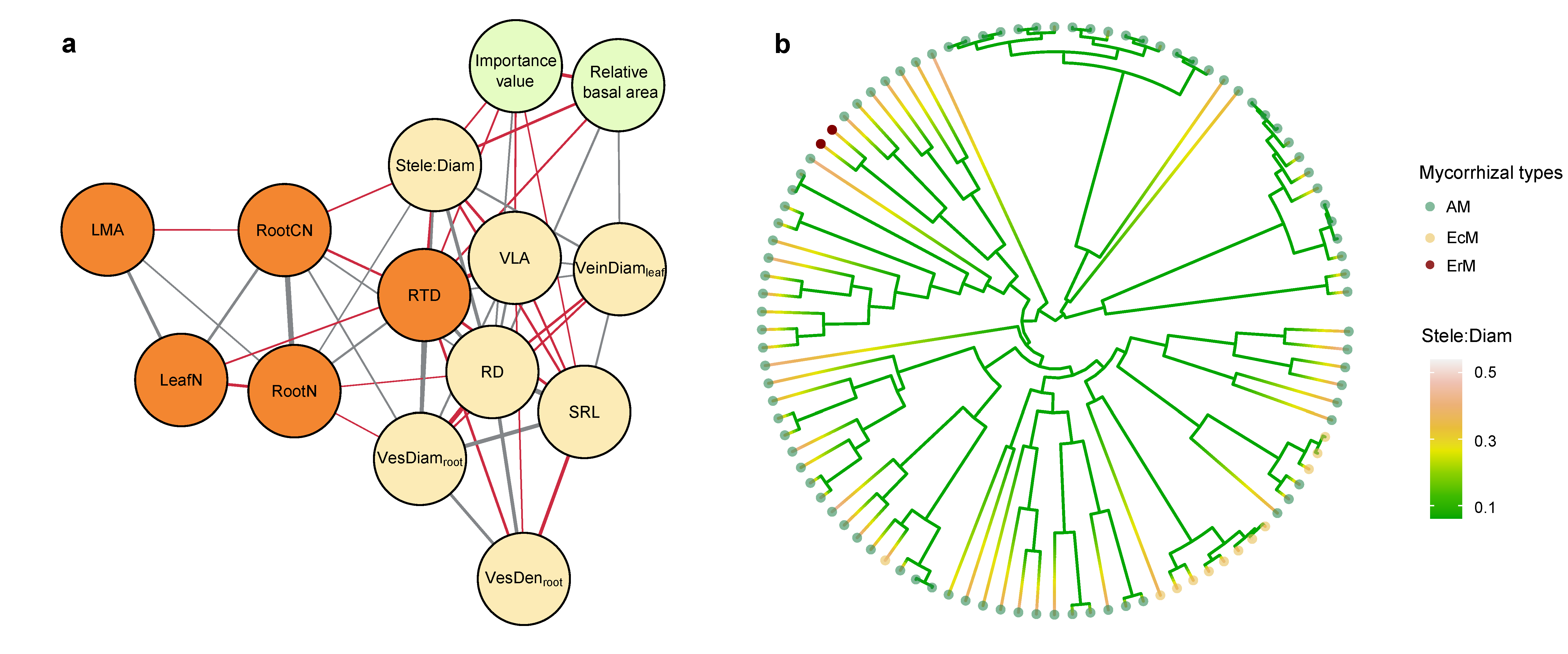

**Extended Data Figure 1 | Trait networks suggest flux-related root-leaf traits link to the species’ importance value (IV). a,** Root and leaf trait networks quantify the correlative structure and functions of whole plants, reflecting the role of plant organisms within the community. Root stele diameter ratios (Stele:Diam), vessel diameter (VesDiam_root_), and leaf nitrogen concentrations (LeafN) are correlated with importance value of species. Leaf minor vein density (VLA) and diameter (VeinDiam_leaf_) are linked with root hydraulic flux traits, particularly root diameter (RD), root vessel diameter (VesDiam_root_), and vessel density (VesDens_root_). **b,** we reconstructed species phylogeny by determining the genomie DNA of each species of plant leaves. Stele:Diam is evolutionary conserved (Pagel’s λ = 0.42, *P* < 0.001), with younger species possessing higher Stele:Diam values.

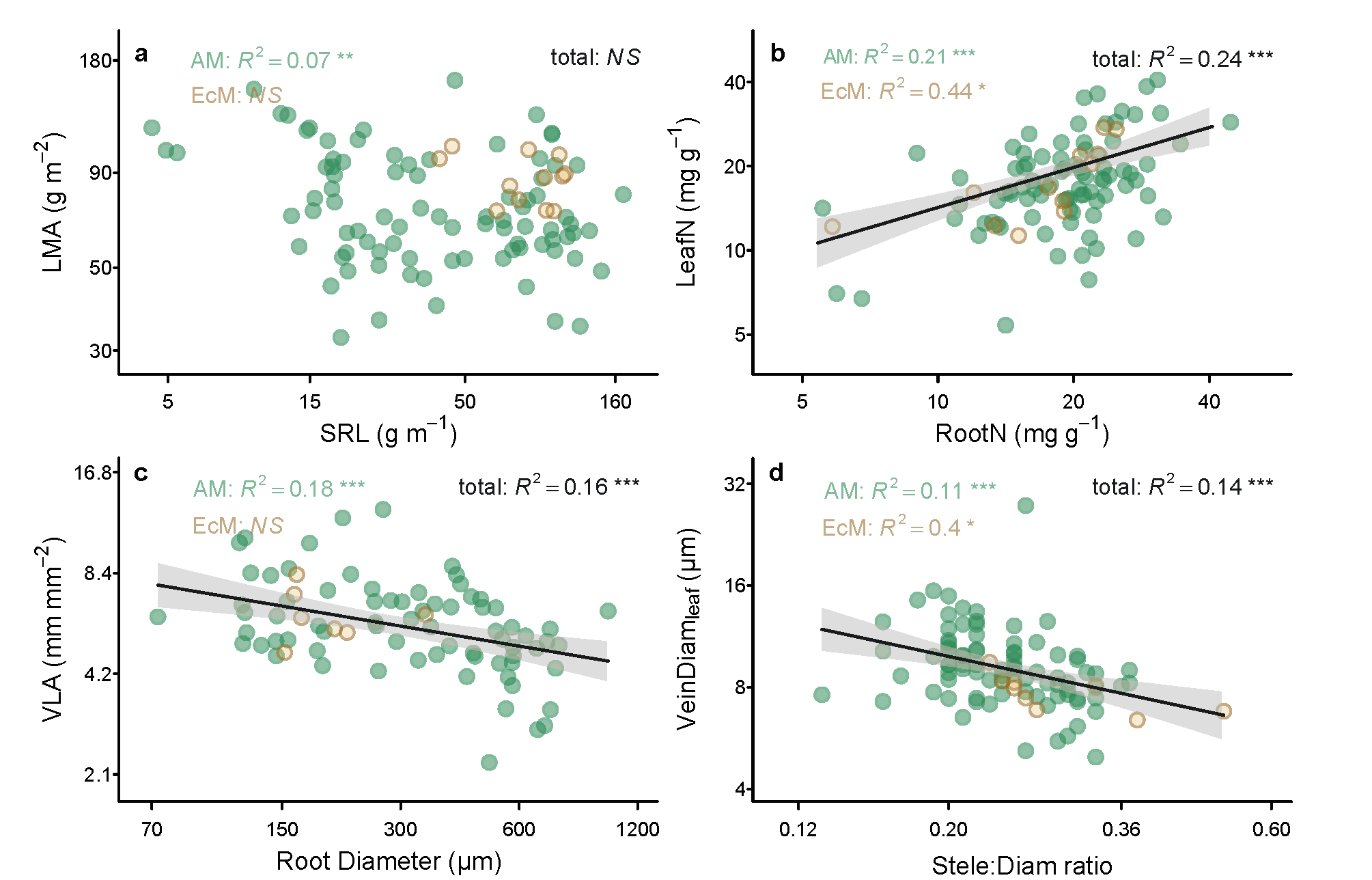

**Extended Data Figure 2 | Root-leaf trait relationships across 101 (sub)-tropical forest species. a,** there is no correlation between leaf mass per area (LMA) and specific root length (SRL) across 101 species (R^2^ = 0.02, *p*=0.07, N=101), while there is a weak correlation between AM species. **b,** there is a positive correlation between LeafN and RootN (R^2^ = 0.24, *p* < 0.001, N=101). **c,** leaf vein density is negatively correlated with root diameter (R^2^ = 0.16, *p* < 0.001, N=82). **d,** the leaf minor vein diameter is negatively correlated with root Stele:Diam (R^2^ = 0.16, *p* < 0.001, n = 96). Each point represents one species. The green dots indicate AM species, and the yellow dots indicate EcM species.

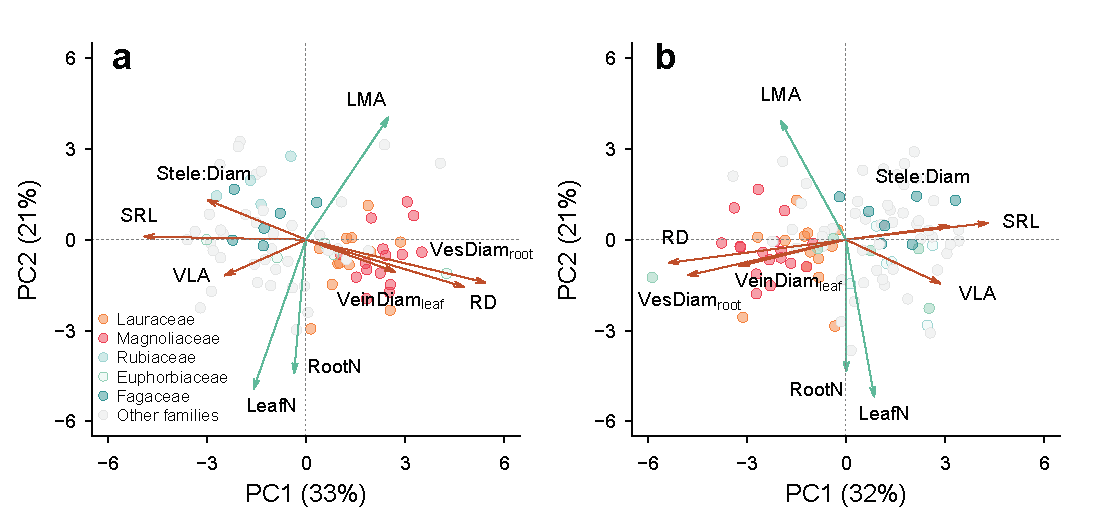

**Extended Data Figure 3 | Phylogenetically informed principal component analysis (pPCA) on root-leaf trait relationships using different datasets. a,** our pPCA using the complete dataset with 82 species revealed two independent axes of the flux-related root-leaf traits (root diameter (RD), specific root length (SRL), root vessel diameter (VesDiam_root_), ratio of root stele to diameter (Stele:Diam), leaf vein density (Wright *et al.*) and minor vein diameter (VeinDiam_leaf_) and economic traits (Leaf mass per area (LMA), Leaf nitrogen concentration (LeafN) and root nitrogen concentration (RootN). **b,** the results of corresponding PCA with the imputed dataset (101 species) are similar to the complete dataset. Each point represents one species. Details are shown in Extended Data Table 6.

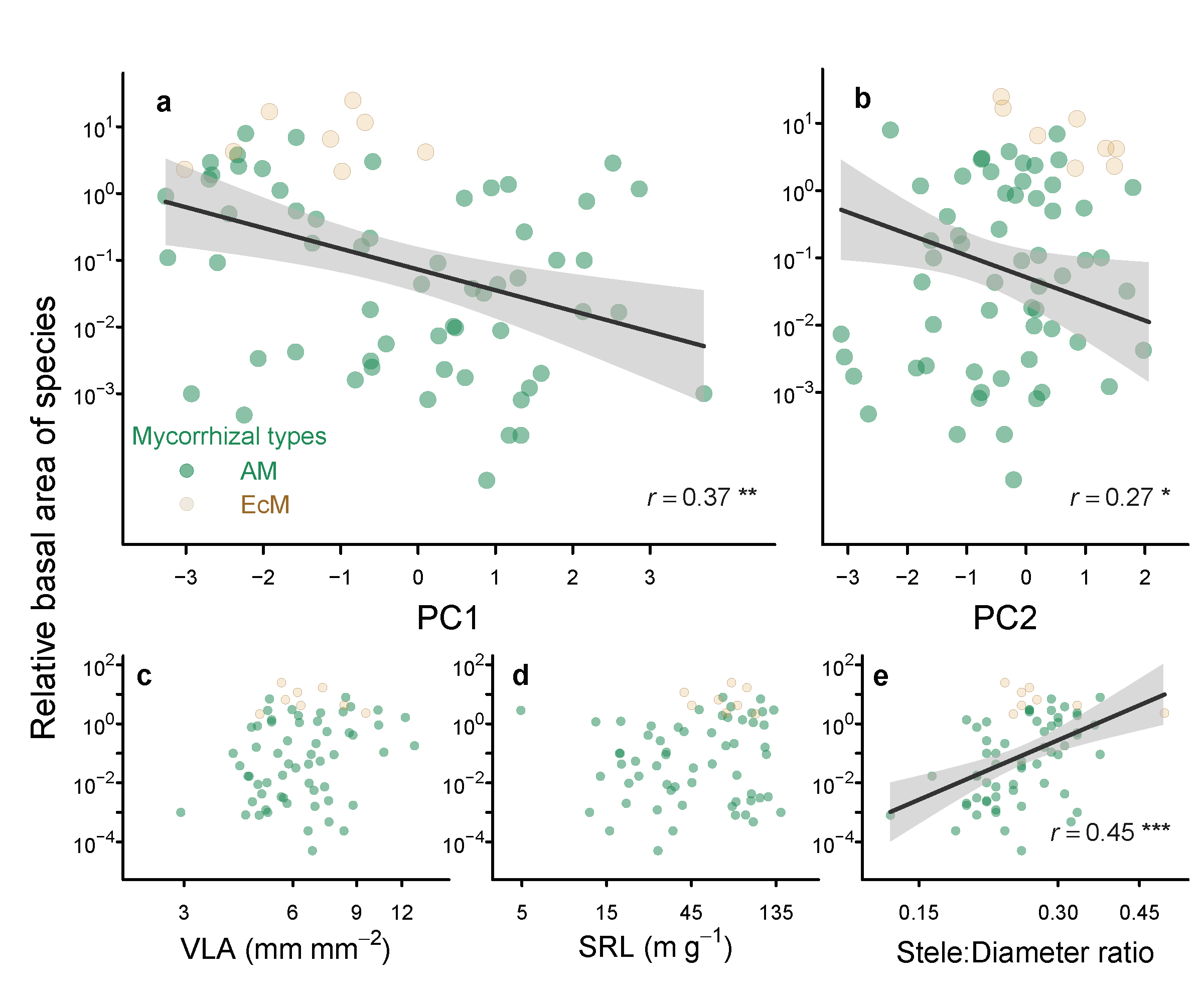

**Extended Data Figure 4 | Flux-related root-leaf traits link to the relative basal area (RBA) across 68 species.** panel **a** showing the correlation between RBA and the PC1 axis, which captures variation in flux-related traits. Species with higher RBA tend to exhibit thinner root, lower leaf vein diameter but greater root stele diameter ratio and vein leaf per leaf area (VLA). **b**, we also found correlation between PC2 and RBA. PC2 is a linear combination of leaf mass per unit area (LMA), leaf nitrogen concentration (LeafN), and root nitrogen concentration (RootN). While RBA does not show a strong correlation with traits (VLA, SRL) related to exchange surface area (**c, d**), it is significantly correlated with Stele: Diameter ratio (**e**), highlighting its importance in water transport efficiency. Yellow and green represent the mycorrhizal fungi type; the EcM species tend to competitively dominate in seasonal forests. The x and y-axis was log scaled.

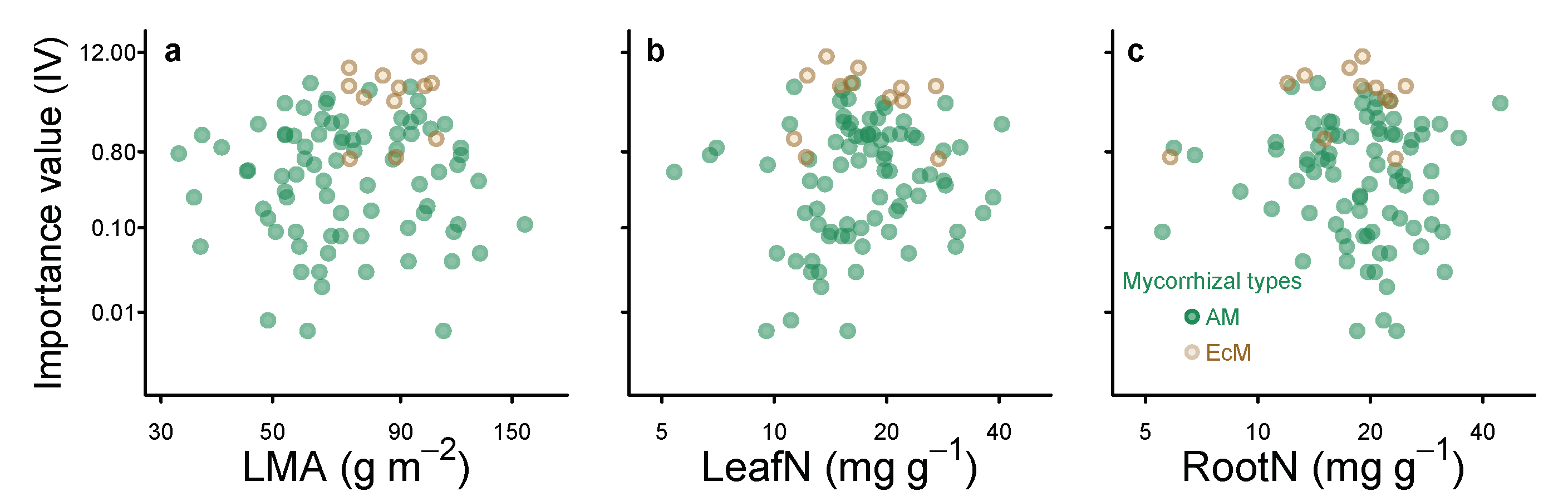

**Extended Data Figure 5 | Root-leaf economic traits (PC2) are not related to the species importance value.** No correlation between PC2 and species importance value. PC2 is a linear combination of leaf mass per tree (LMA), leaf nitrogen concentration (LeafN), and root nitrogen concentration (RootN). **a-c,** the LMA, LeafN, and RootN are not related to the importance values of the species. Yellow and green colors represent species across the mycorrhizal associations.

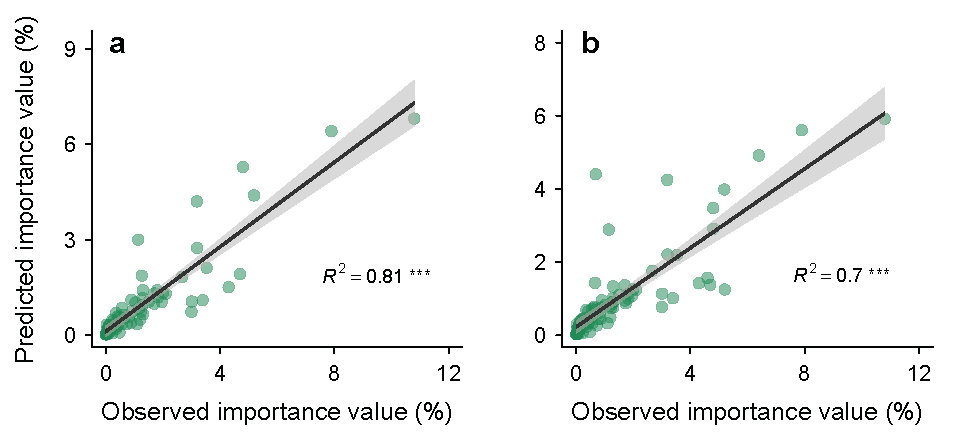

**Extended Data Figure 6 | The prediction of species’ importance value using linear mixed-effect models. a,** the optimal linear mixed-effect model using a complete dataset with 82 species can predict the observed importance value well (R^2^ = 0.81, *p <* 0.001). **b,** the results of the optimal model for the imputed dataset (101 species) are similar to the complete raw dataset (82 species) (R^2^ = 0.70, *p <* 0.001).

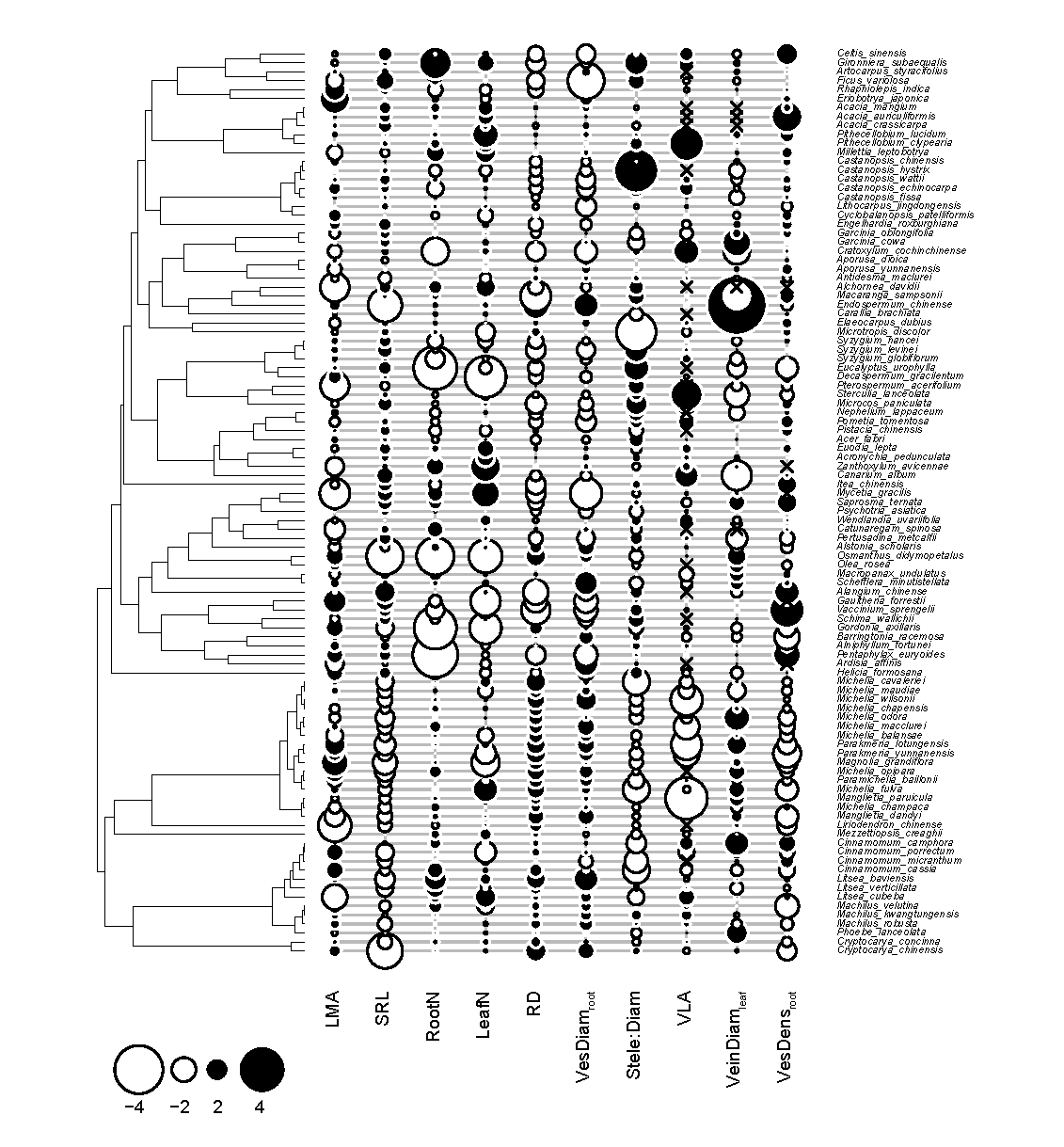

**Extended Data Figure 7 | Phylogeny and relativized trait values mapped onto each of the 101 woody species in this study.** Trait values were centered at zero to facilitate illustration. Black circles represent high values (magnitude is scaled by circle diameter), and white circles represent low values. LMA, leaf mass per area; SRL, specific root length; RootN, root nitrogen concentration; LeafN, leaf nitrogen concentration; VesDiam_root_, root vessel diameter; Stele:Diam, ratio of root stele to diameter; VLA, leaf vein density; VeinDiam_leaf_, leaf minor vein diameter; RD, root diameter; VesDens_root_, root vessel density.

**Extended Data Table 1. Characteristics of the six (sub)-tropical sampling sites in South China.**

| Sites | Longitude | Latitude | Altitude (m) | MAT* (ºC) | MAP^†^ (mm) | Forest type |
| --- | --- | --- | --- | --- | --- | --- |
| Mt. Jianfeng | 108°49´E | 18°40´N | 600-700 | 19.7 | 2651 | Tropical seasonal rain forest |
| Xishuangbanna | 101°25´E | 21°41´N | 500-600 | 21.5 | 1539 | Tropical seasonal rain forest |
| Mt. Wutong | 114°10´E | 22°35´N | 70-100 | 22.4 | 1948 | Subtropical evergreen broadleaved forest |
| Mt. Dinghu | 112°35´E | 23°08´N | 200-300 | 21.4 | 1900 | Subtropical evergreen broadleaved forest |
| Mt. Ailao | 101°01´E | 24°32´N | 2450-2500 | 10.7 | 1841 | Subtropical evergreen broadleaved forest |
| Mt. He | 112°50´E | 22°40´N | 70-90 | 21.7 | 1700 | Subtropical evergreen broadleaved forest |

*MAT = mean annual temperature

^†^MAP = mean annual precipitation

**Extended Data Table 2. Primer sequences used in this study.**

| Primer | Primer sequences |
| --- | --- |
| *rbcL* |  |
| F |  |
| rbc*L* 1F | 5’-ATGTCACCACAAACAGAAAC-3’ |
| rbc*L*a_f | 5’-ATGTCACCACAAACAGAGACTAAAGC-3’ |
| R |  |
| rbcL 724R | 5’-TCGCATGTACCTGCAGTAGC-3’ |
| *matK* |  |
| F |  |
| matK 1R_KIM | 5’-ACCCAGTCCATCTGGAAATCTTGGTTC-3’ |
| matK 390F | 5’-CGATCTATTCATTCAATATTTC-3’ |
| xf | 5’-TAATTTACGATCAATTCATTC-3’ |
| R |  |
| matK 3F_KIM | 5’-CGTACAGTACTTTTGTGTTTACGAG-3’ |
| matK 1326R | 5’-TCTAGCACACGAAAGTCGAAGT-3’ |
| 5r | 5’-GTTCTAGCACAAGAAAGTCG-3’ |

**Extended Data Table 3.** **Fossil information, prior distribution of fossil calibration nodes and the estimated divergence time.**

| Node | MRCA | Stratigraphic position | Age (Mya) | Fossil | Gene fragment | Mean  (Mya) | 95%HPD  (Mya) | Prior distribution |
| --- | --- | --- | --- | --- | --- | --- | --- | --- |
| 1 | Nymphaeales | Late Aptian | [125.0,112.0] | *Endressinia brasiliana* small peryginous flower | *rbcL*  *matK*  *rbcL+matK* | 113.16  112.12  112.03 | [105.64,119.74]  [105.35,118.86]  [105.61,118.64] | Normal  Mean=118,  SD=4 |
| 2 | Magnoliales | Late Aptian | [125.0,112.0] | *Endressinia brasiliana* leaves and flowers | *rbcL*  *matK*  *rbcL+matK* | 115.63  115.68  115.60 | [106.69,123.06]  [108.84,122.32]  [108.54,122.48] | Normal  Mean=118,  SD=4 |
| 3 | Lauraceae | Early Albian-mid Albian | [112.0,99.6] | *Potomacanthus lobatus* flower | *rbcL*  *matK*  *rbcL+matK* | 104.84  104.61  104.68 | [97.63,112.69]  [97.45,111.94]  [97.46,112.36] | Normal  Mean=106,  SD=4 |
| 4 | Magnoliaceae | Early Cenomanian | [99.6,93.5] | *Archaeanthus linnenbergeri* fruit | *rbcL*  *matK*  *rbcL+matK* | 96.47  96.37  96.42 | [92.60,100.50]  [92.64,100.19]  [92.48,100.35] | Normal  Mean=96.5,  SD=2 |
| 5 | Fagales | Late Cenomanian | [96.5,93.5] | *Normapolles* pollen | *rbcL*  *matK*  *rbcL+matK* | 94.30  94.19  94.19 | [93.50,95.75]  [93.50,95.57]  [93.50,95.50] | exponential  Offset=93.5,  Mean=1 |
| 6 | Ericales | Turonian | [93.5,89.3] | *Paleoenkianthus sayrevillensis* flowers | *rbcL*  *matK*  *rbcL+matK* | 91.22  91.15  91.12 | [88.35,94.11]  [88.03,93.89]  [88.21,93.98] | Normal  Mean=91.5,  SD=1.5 |
| 7 | Malpighiales | Turonian | [93.5,89.3] | *Paleoclusia chevalieri* flowers | *rbcL*  *matK*  *rbcL+matK* | 91.31  90.91  90.95 | [88.48,94.40]  [88.21,93.79]  [88.15,93.91] | Normal  Mean=91.5,  SD=1.5 |
| 8 | Sapindales | Late Paleocene | [58.7,55.8] | *Acer* sp., *Dipteronia* sp., *Koelreuteria* sp. fruits and leaves | *rbcL*  *matK*  *rbcL+matK* | 57.14  57.15  57.17 | [55.22,59.05]  [55.12,59.04]  [55.24,59.15] | Normal  Mean=57  SD=1 |
| 9 | Rosaceae | Middle Eocene | [48.6,37.2] | *Prunus* sp. endocarp, *Rosa* sp. foliage | *rbcL*  *matK*  *rbcL+matK* | 41.94  42.05  42.14 | [36.29,47.95]  [36.23,47.73]  [36.42,47.93] | Normal  Mean=43  SD=3 |
| 10 | Malvales | Late Eocene | [37.2,33.9] | *Craigia* sp., *Tilia* sp. flowers | *rbcL*  *matK*  *rbcL+matK* | 35.47  35.48  35.48 | [33.59,37.43]  [33.46,37.37]  [33.50,37.36] | Normal  Mean=35.5,  SD=1 |
| 11 | Eudicot | Late Barremian-Early Aptian Barremian-Early aptian | [130,125] | Tricolpate pollen grains *Sinocarpus decussatus* infructescence fragment | *rbcL*  *matK*  *rbcL+matK* | 126.03  126.51  126.50 | [125.00,128.01]  [125.00,129.34]  [125,129.20] | exponential  Offset=125  Mean=1.5 |
| 12 | Rutaceae | Late Eocene | [37.2,33.9] | *Euodia* sp. seeds, *Rutaspermum* sp. seeds, *Ptelea* sp. fruit | *rbcL*  *matK*  *rbcL+matK* | 35.31  35.29  35.26 | [33.44,37.31]  [33.45,37.37]  [33.36,37.26] | Normail  Mean=35.5,  SD=1 |

HPD, Highest posterior density.

Fossil age refer to GTS 2004 (Gradstein F, Ogg J. 2004. Geologic time scale 2004 – why, how, and where next! *Lethaia* 37: 175-181.)

**Extended Data Table 4. Coefficients of Spearman correlations for pairwise traits with complete dataset (lower-left diagonal) and phylogenetically independent contrasts (upper-right diagonal).**

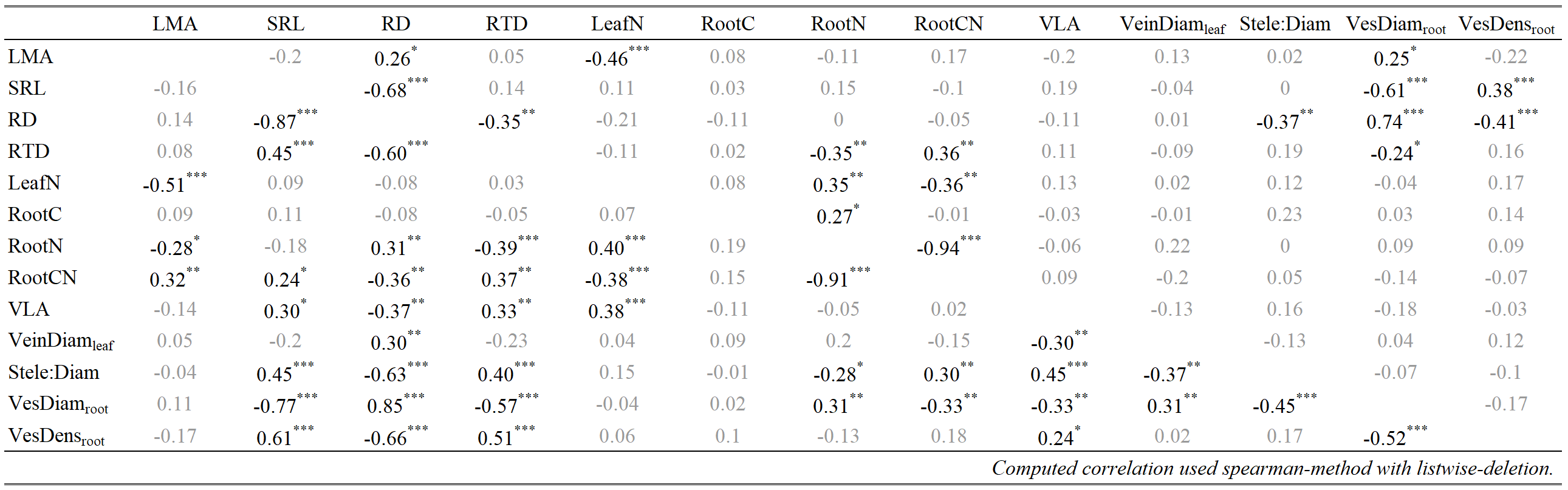

Significant correlations are indicated; * * * P < 0.001; * * P < 0.01; * P < 0.05 and are shown in black text while non-significant correlations are in gray text.

**Extended Data Table 5. Coefficients of Spearman's correlation for pairwise traits with imputed dataset (lower-left diagonal) and phylogenetically independent contrasts (upper-right diagonal).**

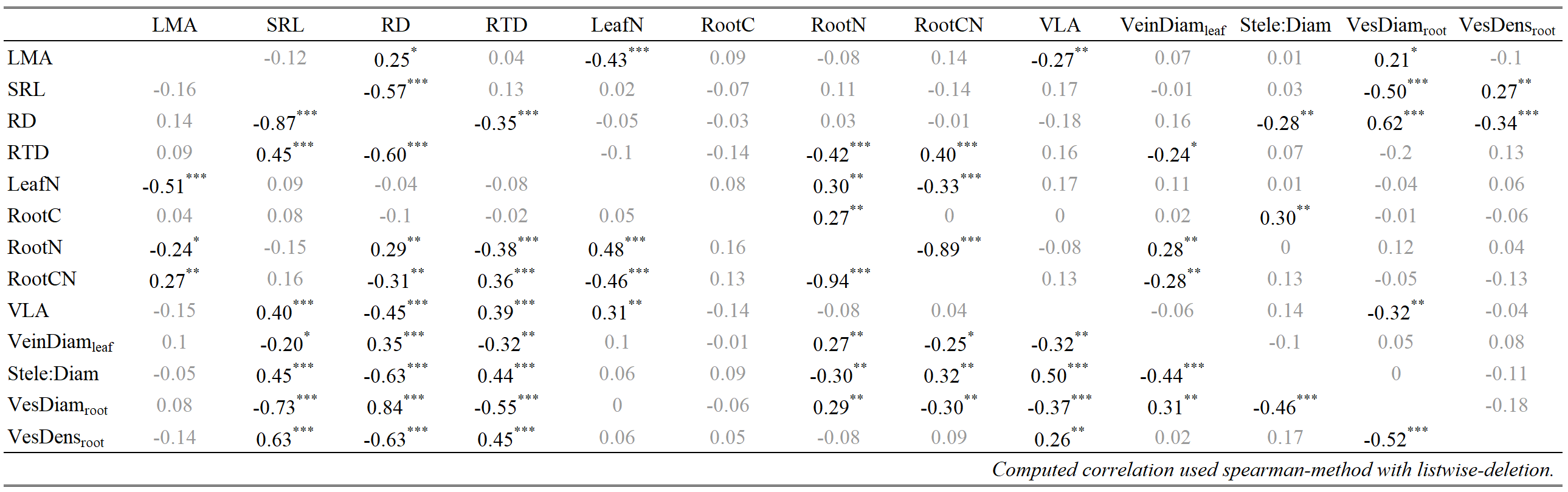

Significant correlations are indicated; * * * P < 0.001; * * P < 0.01; * P < 0.05 and are shown in black text while non-significant correlations are in gray text.

**Extended Data Table 6. Loading scores of absorptive root and leaf traits on principal components 1 and 2 of the unrotated principal components analysis (PCA), the varimax rotated PCA and the phylogenetic PCA for both the imputed dataset (N = 101) and the complete dataset (N = 82).**

| Trait | Imputed dataset (N = 101) | | | | | |  | Complete dataset (N = 82) | | | | | |
| --- | --- | --- | --- | --- | --- | --- | --- | --- | --- | --- | --- | --- | --- |
|  | unrotated PCA | | varimax PCA | | phylogenetic PCA | |  | unrotated PCA | | varimax PCA | | phylogenetic PCA | |
|  | PC1 | PC2 | PC1 | PC2 | PC1 | PC2 |  | PC1 | PC2 | PC1 | PC2 | PC1 | PC2 |
| LMA | -0.25 | **0.70** | 0.20 | **-0.71** | -0.33 | **0.66** |  | -0.24 | **-0.73** | 0.18 | **-0.75** | -0.42 | **0.67** |
| SRL | **0.80** | 0.02 | **-0.79** | 0.04 | **0.72** | 0.09 |  | **0.83** | -0.08 | **-0.83** | -0.01 | **0.82** | 0.02 |
| RD | **-0.92** | -0.04 | **0.92** | -0.03 | **-0.90** | -0.13 |  | **-0.93** | 0.05 | **0.94** | -0.02 | **-0.91** | -0.24 |
| RootN | -0.23 | **-0.74** | 0.28 | **0.72** | 0.00 | **-0.73** |  | -0.20 | **0.74** | 0.25 | **0.72** | 0.06 | **-0.73** |
| LeafN | 0.11 | **-0.88** | -0.04 | **0.89** | 0.14 | **-0.86** |  | 0.11 | **0.87** | -0.04 | **0.87** | 0.26 | **-0.82** |
| VLA | **0.64** | -0.26 | **-0.62** | 0.30 | **0.48** | -0.24 |  | **0.58** | 0.31 | **-0.56** | 0.35 | **0.41** | -0.20 |
| VeinDiam_leaf_ | **-0.55** | -0.10 | **0.55** | 0.06 | **-0.54** | -0.15 |  | **-0.52** | 0.04 | **0.52** | 0.00 | **-0.45** | -0.18 |
| VesDiam_root_ | **-0.84** | -0.10 | **0.85** | 0.04 | **-0.80** | -0.20 |  | **-0.87** | 0.13 | **0.87** | 0.06 | **-0.80** | -0.26 |
| Stele:Diam | **0.69** | 0.11 | **-0.69** | -0.06 | **0.52** | 0.08 |  | **0.70** | -0.08 | **-0.70** | -0.02 | **0.50** | 0.22 |
| Variation explained (%) | 39 | 21 | 39 | 21 | 32 | 21 |  | 39 | 22 | 39 | 22 | 33 | 21 |

Eigenvalues and loading scores of principal components 1 and 2 (PC1 and PC2) in three different principal component analyses. For all three analyses, the loading scores show that the PC1 axis is composed of RD, SRL, VLA, VeinDiam_leaf_, VesDiam_root_, stele:Diam, and the PC2 axis is mainly influenced by LMA, LeafN, and RootN. Loadings considered statistically significant (P < 0.05) are shown in black text while non-significant loadings are in gray text.

**Extended Data Table 7. Standardized major axis regression analyses of pairwise relationships across AM and EcM species respectively.**

| Pair of traits | MYT | Slope | Elevation | R^2^ | *p* | N | slope homogeneity (p) |
| --- | --- | --- | --- | --- | --- | --- | --- |
| LMA-SRL | AM | -0.41 | 2.51 | 0.08 | 0.01 | 87 | 0.74 |
|  | EcM | -0.46 | 2.80 | 0.09 | 0.35 | 12 |  |
| LeafN-RootN | AM | 1.06 | -0.12 | 0.22 | **<0.001** | 87 | 0.20 |
|  | EcM | 0.77 | 0.29 | 0.49 | **0.01** | 12 |  |
| VLA-Stele:Diam | AM | 1.51 | 1.70 | 0.20 | **<0.001** | 73 | 0.99 |
|  | EcM | 1.50 | 1.68 | 0.63 | **0.03** | 7 |  |
| VLA-SRL | AM | 0.38 | 0.18 | 0.15 | <0.001 | 73 | 0.55 |
|  | EcM | -0.50 | 1.72 | 0.00 | 0.94 | 7 |  |
| VLA-VesDiam_root_ | AM | -0.77 | 1.48 | 0.14 | 0.00 | 73 | 0.89 |
|  | EcM | -0.73 | 1.34 | 0.08 | 0.54 | 7 |  |
| VLA-VeinDiam_leaf_ | AM | -1.18 | 1.90 | 0.09 | **0.01** | 73 | 0.41 |
|  | EcM | -1.75 | 2.38 | 0.05 | 0.63 | 7 |  |
| VesDiam_root_-VeinDiam_leaf_ | AM | 1.58 | -0.60 | 0.13 | <0.001 | 85 | 0.57 |
|  | EcM | -1.93 | 2.48 | 0.24 | 0.19 | 9 |  |
| VesDiam_root_-SRL | AM | -0.48 | 1.69 | 0.51 | <0.001 | 87 | 0.10 |
|  | EcM | 0.84 | -0.80 | 0.00 | 0.90 | 12 |  |
| VeinDiam_leaf_-Stele:Diam | AM | -1.22 | 0.21 | 0.12 | **0.00** | 85 | 0.01 |
|  | EcM | -0.47 | 0.63 | 0.48 | **0.04** | 4 |  |

**Extended Data Table 8. Patterns of dissimilarity on nine leaf-root traits across different functional groups (mycorrhizal type, leaf habit and growth form).**

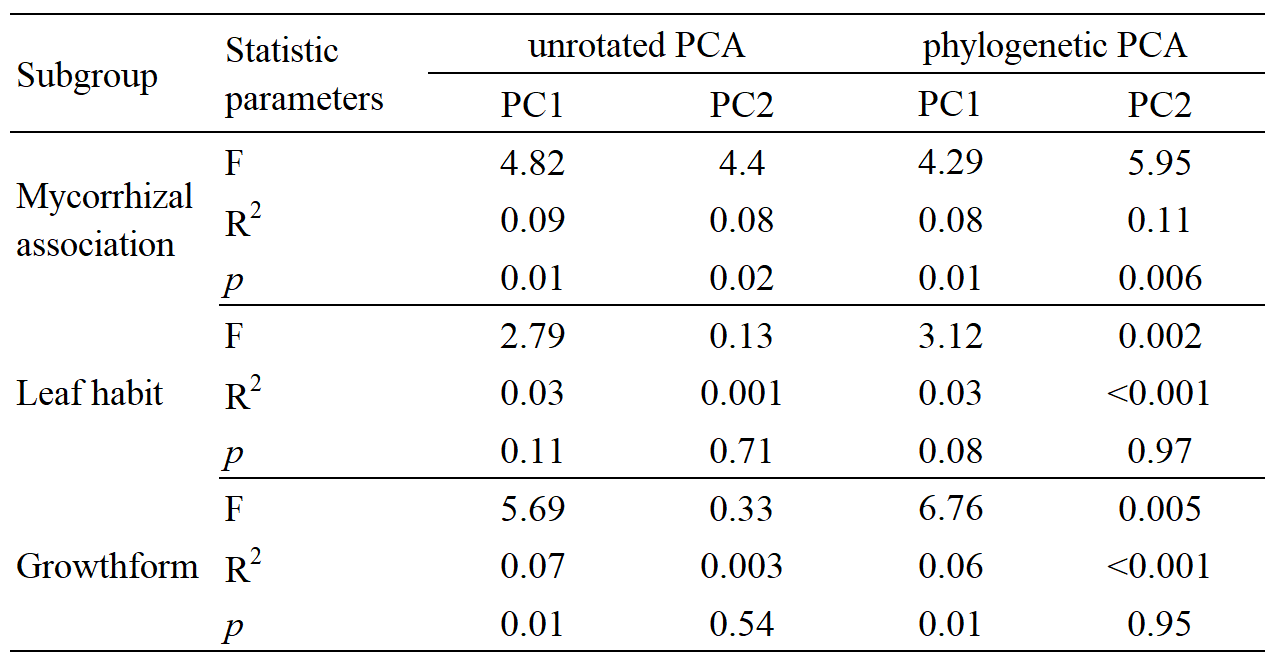

**Extended Data Table 9. Comparison of linear mixed models for the complete dataset (N = 82) and imputed dataset (N = 95) estimating the effect of mycorrhizal types (MYT), root diameter (RD), Vessel diameter (VesDiam_root_), specific root length (SRL), leaf vein density (VLA), and leaf nitrogen (LeafN) on species importance value.**

| Dataset | Step | Model | n par | χ^2^ | p-value | R^2^m | R^2^c | AIC | AIC weight |
| --- | --- | --- | --- | --- | --- | --- | --- | --- | --- |
| Complete dataset  (N = 82) | 0 | Intercept + (1\|Sites) + (1\|Genus) + (1\|Species) | 5 |  |  | 0.00 | 0.56 | 154.5 | 0.000 |
|  | 1 | MYT + (1\|Sites) + (1\|Genus) + (1\|Species) | 6 | 7.00 | 0.008 | 0.18 | 0.54 | 149.5 | 0.002 |
|  | 2 | MYT + RD + (1\|Sites) + (1\|Genus) + (1\|Species) | 7 | 0.55 | 0.460 | 0.20 | 0.54 | 150.9 | 0.001 |
|  | 3 | MYT + RD + VesDiam_root_ + (1\|Sites) + (1\|Genus) + (1\|Species) | 8 | 12.16 | 0.000 | 0.29 | 0.65 | 140.8 | 0.191 |
|  | 4 | MYT + RD + VesDiam_root_ + VLA + (1\|Sites) + (1\|Genus) + (1\|Species) | 9 | 2.62 | 0.106 | 0.35 | 0.70 | 140.2 | 0.261 |
|  | **5** | **MYT + RD + VLA + VesDiam_root_ + LeafN + (1\|Sites) + (1\|Genus) + (1\|Species)** | **10** | **2.71** | **0.100** | **0.36** | **0.76** | **139.4** | **0.371** |
|  | 6 | MYT + RD + LeafN + VLA + VesDiam_root_ + SRL + (1\|Sites) + (1\|Genus) + (1\|Species) | 11 | 0.48 | 0.489 | 0.37 | 0.76 | 141.0 | 0.173 |
| Imputed dataset  (N = 101) | 0 | Intercept + (1\|Sites) + (1\|Genus) + (1\|Species) | 5 |  |  | 0.00 | 0.56 | 187.6 | 0.001 |
|  | 1 | MYT + (1\|Sites) + (1\|Genus) + (1\|Species) | 6 | 8.67 | 0.003 | 0.16 | 0.53 | 180.9 | 0.021 |
|  | 2 | MYT + RD + (1\|Sites) + (1\|Genus) + (1\|Species) | 7 | 0.22 | 0.637 | 0.17 | 0.53 | 182.7 | 0.009 |
|  | 3 | MYT + RD + VesDiam_root_ + (1\|Sites) + (1\|Genus) + (1\|Species) | 8 | 8.06 | 0.005 | 0.20 | 0.64 | 176.6 | 0.181 |
|  | 4 | MYT + RD + VesDiam_root_ + VLA + (1\|Sites) + (1\|Genus) + (1\|Species) | 9 | 2.75 | 0.097 | 0.26 | 0.66 | 175.9 | 0.264 |
|  | **5** | **MYT + RD + VLA + VesDiam_root_ + LeafN + (1\|Sites) + (1\|Genus) + (1\|Species)** | **10** | **2.54** | **0.111** | **0.27** | **0.72** | **175.3** | **0.346** |
|  | 6 | MYT + RD + LeafN + VLA + VesDiam_root_ + SRL + (1\|Sites) + (1\|Genus) + (1\|Species) | 11 | 0.66 | 0.415 | 0.27 | 0.72 | 176.7 | 0.178 |

We used a linear mixed-effect model to evaluate the impact of different root-leaf traits on community importance value (IV), with species phylogeny and sampling sites as the random effect. The marginal R^2^ (R^2^m) describes the goodness of model fit given fixed effects only, while the conditional R^2^ (R^2^c) describes the goodness of model fit including fixed and random effects.
